# NeurostimML: A machine learning model for predicting neurostimulation-induced tissue damage

**DOI:** 10.1101/2023.10.18.562980

**Authors:** Yi Li, Rebecca A. Frederick, Daniel George, Stuart F. Cogan, Joseph J. Pancrazio, Leonidas Bleris, Ana G. Hernandez-Reynoso

**Affiliations:** Department of Bioengineering, The University of Texas at Dallas, Richardson, TX, USA; Center for Systems Biology, The University of Texas at Dallas, Richardson, TX, USA; Phil and Penny Knight Campus for Accelerating Scientific Impact, University of Oregon, Eugene, OR, USA; Department of Computer Science, The University of Texas at Dallas, Richardson, TX, USA; Department of Biological Sciences, The University of Texas at Dallas, Richardson, TX, USA

**Keywords:** neuromodulation, Shannon equation, machine learning, safe stimulation

## Abstract

**Objective:** The safe delivery of electrical current to neural tissue depends on many factors, yet previous methods for predicting tissue damage rely on only a few stimulation parameters. Here, we report the development of a machine learning approach that could lead to a more reliable method for predicting electrical stimulation-induced tissue damage by incorporating additional stimulation parameters.

**Approach:** A literature search was conducted to build an initial database of tissue response information after electrical stimulation, categorized as either damaging or non-damaging. Subsequently, we used ordinal encoding and random forest for feature selection, and investigated four machine learning models for classification: Logistic Regression, K-nearest Neighbor, Random Forest, and Multilayer Perceptron. Finally, we compared the results of these models against the accuracy of the Shannon equation.

**Main Results:** We compiled a database with 387 unique stimulation parameter combinations collected from 58 independent studies conducted over a period of 47 years, with 195 (51%) categorized as non-damaging and 190 (49%) categorized as damaging. The features selected for building our model with a Random Forest algorithm were: waveform shape, geometric surface area, pulse width, frequency, pulse amplitude, charge per phase, charge density, current density, duty cycle, daily stimulation duration, daily number of pulses delivered, and daily accumulated charge. The Shannon equation yielded an accuracy of 63.9% using a k value of 1.79. In contrast, the Random Forest algorithm was able to robustly predict whether a set of stimulation parameters was classified as damaging or non-damaging with an accuracy of 88.3%.

**Significance:** This novel Random Forest model can facilitate more informed decision making in the selection of neuromodulation parameters for both research studies and clinical practice. This study represents the first approach to use machine learning in the prediction of stimulation-induced neural tissue damage, and lays the groundwork for neurostimulation driven by machine learning models.

## 1. Introduction

Neuromodulation of the central and peripheral nervous systems has been widely used to restore function after injury or disease. Generally, neuromodulation by electrical stimulation is used as a last line of treatment for refractory or intractable conditions that have no other alternatives to achieve relief or restore function. Some of the most successful examples of neuromodulation include deep brain stimulation (DBS) to treat Parkinson’s disease and essential tremor^1^, cochlear implants to restore hearing^2^, vagus nerve stimulation to treat epilepsy^3^, reduce inflammation^4^, and to improve motor function after stroke or spinal cord injury when paired with rehabilitation^5,6^, sacral nerve stimulation to treat overactive bladder and fecal incontinence^7,8^, and spinal cord stimulation to treat paralysis and chronic pain^9,10^. Despite the many successful implementations of electrical neuromodulation to treat disease and restore function, the potential damage that electrical stimulation can cause to neural tissues remains a concern. The evaluation of neural stimulation safety relies on only a limited set of parameters derived from a set of important yet limited systematic studies^11–17^. Much of the reason for the inability to precisly define when electrical stimulation-induced tissue damage will occur lies in the complexity of processes occurring during neuromodulation, and the large number of stimulation waveform parameters available to modulate in addition to electrochemical and physiological variables. In addition, electrode selection and design can influence electric field distribution in the tissue, result in non-reversible charge-transfer reactions at the electrode-tissue interface, and electrode dissolution, which may also lead to stimulation-induced tissue damage.

Several experimental studies have systematically investigated the relationship between stimulation parameters and the potential for tissue damage^12,18,19^. However, the development of predictive models for neural tissue damage has been hindered by the relatively limited availability of experimental data when compared to the number of variables involved.

The most well-known and accepted mathematical relationship for predicting stimulation-induced tissue damage is the Shannon equation^20^ that was published in 1992 “as an initial framework for designing future animal safety studies”. The model describes a logarithmic two-dimensional mathematical boundary to separate likely damaging from likely non-damaging stimulation parameters based on data from stimulation damage observed in cat cortex using macroelectrodes (GSA = 0.01 to 0.5 cm^2^, which are much larger than the GSA ranges for common microelectrodes, typically ranging between 200 - 2000 µm^2^). The equation relates the charge per phase and the charge density (charge per unit electrode surface, delivered within the leading phase of a charge-balanced stimulus pulse) (**Equation 1**). However, the Shannon model is based on only “near-field” histology data and does not account for any additional parameters beyond charge per phase and charge density per phase that may contribute to stimulation-induced damage. Shannon himself states in the seminal 1992 publication that “A more comprehensive model of safe levels for electrical stimulation would also account for the effects of pulse rate, pulse duration, stimulus duty cycle, and duration of exposure.”^20^ There are a number of neuromodulation cases requiring descriptions outside the capabilities of the Shannon model, and a more detailed discussion of these cases can be found in Cogan et al.^21^.

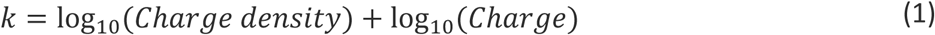

Vatsyayan and Dayeh^22^ expanded the Shannon model in 2022 to account for the electrode site material, electrode size, and inter-electrode spacing. However, their model focuses primarily on electrochemical reactions and still does not incorporate other parameters that have been shown to play a role in the design of safe stimulation protocols. In particular, neither the Shannon nor Vatsyayan-Dayeh models incorporate stimulation parameters related to the cumulative nature or time scale of charge delivery (e.g., frequency or duty cycle). Each model does, however, point to the complexity of the relationship between stimulation parameters and tissue damage. Even within the two-parameter space Shannon model, the features indicating tissue damage exhibit non-linearity as the identification of a boundary between damaging and non-damaging stimulation requires comparing the log_10_ values of both charge per phase and charge density per phase. The Vatsyayan-Dayeh model describes the complex interactions between the non-linear features of electrochemical impedance and charge injection capacity, which are related to material-dependent charge injection mechanisms that can change with stimulation frequency and amplitude.

Supervised and unsupervised machine learning has revolutionized diverse research fields from social sciences to biological sciences and medicine^23–28^. In supervised learning (e.g., k-nearest neighbours or multilayer perceptron), the non-linear relationship between input variables (features) and output targets (labels) is uncovered using training instances, which can be subsequently used for prediction on new instances (testing instances)^29^. Supervised learning algorithms have been widely used in areas of neuroscience and neural engineering research including brain imaging analysis^30–32^, neuroinformatics^33–35^, and behavioural analysis^36–38^.

Herein, we develop a machine learning model to predict electrical stimulation-induced neural tissue damage. Our primary objective was to explore the utility of a machine learning to create a more effective tool for estimating safe thresholds for electrical neuromodulation parameters. By utilizing published experimental data and machine learning methods to identify stimulation parameters indicative of tissue damage, we hope to guide the design of additional studies for more targeted investigation of the available parameter space and the underlying mechanisms leading to stimulation-induced tissue damage.

## 2. Materials and Methods

### 2.1 Literature search and data collection

For initial data collection, we first utilized the bibliographic references included in recent review articles addressing safety of neural stimulation: Cogan et al. (2016)^21^, Shepherd et al. (2006)^39^, Miller et al. (2001)^43^, and Agnew et al. (1990)^12^. We then performed a literature search by first generating a comprehensive list of phrases and terms within three key search areas: electrical stimulation of neural tissue, tissue damage, and neural interface device and electrode features. We then identified and removed overlapping or redundant terms and built the final search query in PubMed on February 15, 2023:

> (((“Electric Stimulation” [MeSH Terms] OR “Nerve Stimulation” [Text Word] OR “Brain Stimulation” [Text Word] OR “Spinal Cord Stimulation” [Text Word] OR “Microstimulation” [Text Word] OR “Neuromodulation” [Text Word] OR “Stimulation Parameters” [Text Word] OR “Pulsing” [Text Word])) AND ((“Nerve Degeneration” [MeSH Terms] OR “Wallerian Degeneration” [MeSH Terms] OR “Tissue Assessment” [Text Word] OR “Tissue Response” [Text Word] OR “Histopathology” [Text Word] OR “Tissue Damage” [Text Word] OR “Damage” [Text Word] OR “Damaging” [Text Word])) AND ((“Neural Prostheses” [MeSH Terms] OR “Neural Prosthesis” [Text Word] OR “Electrodes” [MeSH Terms] OR “Microelectrodes” [MeSH Terms] OR “Implantable Neurostimulators” [MeSH Terms] OR “Neural Interface” [Text Word] OR “Microelectrode Array” [Text Word] OR “Electroceutical” [Text Word])) AND ((English [Filter]))) NOT (Review[Publication Type])

Inclusion criteria for the literature search were defined as: (1) studies containing primary sources (e.g., experimental data not review articles), (2) studies reporting both electrical stimulation parameters and histopathological assessment of the stimulated neural tissue. Exclusion criteria for the literature search were defined as: (1) review articles or articles containing secondary or tertiary reports of experimental data, (2) studies using non-invasive stimulation, (3) in vitro studies, (4) retinal stimulation studies, (5) studies combining electrical stimulation with pharmacological agents (other than antibiotics or those used for pain management), (6) studies that do not report any histopathological assessment.

From each of the included studies, we compiled 12 categorical and 19 numerical features, in addition to an indexing feature (Reference ID). The list of all categorical and numerical features, definitions and units can be found in **Table 1**; a visual explanation of stimulation parameters and waveforms is shown in **Supplementary Figure 1**. We included as separate entries all stimulation parameter combinations for articles reporting multiple parameters. For those investigating the same parameters in multiple subjects, we included as a single entry the average outcome for all subjects. All entries were assigned a score to report the histological or functional outcome either from the original assessment reported by the source, or from scoring performed by two investigators (authors RAF and AGH-R) blinded to each other’s assessments. For the latter, investigators graded the outcome on a scale from 0 to 4 corresponding to 0 – no damage, 1 – minimal damage (similar to implanted but unpulsed electrodes), 2 – mild damage, 3 – moderate damage, and 4 – severe damage. Then, binary scores were calculated based on the averaged graded responses between investigators as 0 – non-damaging (if average graded outcome ≤ 1) and 1 – damaging (if average graded outcome > 1).

**Table 1.** List of features (inputs) and labels (outputs) for our current study.

| Feature | Type | Description | Units |
| --- | --- | --- | --- |
| Entry | Numerical | Unique ID number for each data entry. | none |
| Reference ID | Alphanumeric | Citation key to identify reference source. | none |
| Animal | Categorical | Animal model used in cited reference. | none |
| Nervous_system | Categorical | Label describing if stimulation targeted central or peripheral nervous system. | none |
| Tissue_region | Categorical | Anatomical region targeted for stimulation. | none |
| Tissue_target<br>_structure | Categorical | Implant location within the tissue region. | none |
| Circuit_control | Categorical | Label describing if stimulation was voltage-controlled or current-controlled. | none |
| Waveform_shape | Categorical | Shape of the waveform used for stimulation. | none |
| Initial_current<br>_direction | Categorical | Direction of the leading phase in the stimulation pulse. | none |
| Array_configuration | Categorical | Number of 'poles' used for spatial arrangement of stimulation electrodes. | none |
| Electrode_shape | Categorical | Basic geometric shape of the electrode. | none |
| Electrode_material | Categorical | Material composition of the electrode. | none |
| Charge_mechanism | Categorical | Charge transfer mechanism, defined by the stimulation electrode material. | none |
| Electrode_GSA | Numerical | Geometric surface area of the electrode. | $\mu\text{m}^2$ |
| Voltage_dc_bias | Mixed | "None" if no bias strategy was used, or numerical value of DC voltage (vs. reference) maintained between stimulation pulses. | V |
| Pulse_width | Numerical | Duration of the leading phase of the stimulation pulse. | $\mu\text{s}$ |
| Frequency | Numerical | Number of stimulation pulses per second. | Hz |
| Interphase_delay | Numerical | Duration of time between first and second phase of the stimulation pulse. (monophasic stimulation = 0) | $\mu\text{s}$ |
| Current_amplitude | Numerical | Absolute value of the current applied in first phase of the stimulation pulse. | $\mu\text{A}$ |
| Voltage_amplitude | Numerical | Absolute value of the voltage applied in first phase of the stimulation pulse. | mV |
| Charge_per_phase | Numerical | Amount of charge within the first phase of the stimulation pulse. | nC/ph |
| Charge_density | Numerical | Distribution of charge over the electrode surface area within the first phase of the stimulation pulse. | $\mu\text{C/ph/cm}^2$ |
| Current_density | Numerical | Distribution of current over the electrode surface area within the first phase of the stimulation pulse. | $\text{mA/cm}^2$ |
| Duty_cycle | Numerical | Proportion of time pulsing was on per day. | % |
| Stim_time_on | Numerical | Time duration that pulsing was on. | s |
| Stim_time_off | Numerical | Time duration that pulsing was off. | s |
| Stim_time_daily | Numerical | Total time duration pulsing was on per day. | s |
| Days_stim | Numerical | Number of days of stimulation during the study. | days |
| Stim_total_study | Numerical | Total number of hours of stimulation during the study. | h |
| Daily_pulses | Numerical | Number of pulses applied per day. | pulses |
| Total_pulses | Numerical | Number of pulses applied over the entire study. | pulses |
| Daily_accumulated<br>_charge | Numerical | Total amount of charge delivered to the tissue per day. | C |
| Total_accumulated<br>_charge | Numerical | Total amount of charge delivered to the tissue over the entire study. | C |
| Damage_level | Numerical | Integer score of 0, 1, 2, 3, or 4. | none |
| Damage_binary | Binary | Translation of integer scores to binary values, where 0-1 = "0" and 2-4 = "1". | none |
| k | Numerical | Coefficient value calculated using the Shannon equation. | none |

### 2.2 Predictions using the Shannon equation

Two features, charge per phase (unit: nC ph^-1^) and charge density per phase (unit: µC cm^-2^ ph^-1^), were used to calculate the *k*-values related to the Shannon equation (Equation 1). We tested a range of *k* threshold values ranging from *k* = -4 to *k* = 4 to obtain a prediction of stimulation outcomes on tissue. The calculated *k*-values below a given threshold (e.g., *k*=1.85) were classified as non-damaging, and those larger than or equal to the given threshold were classified as damaging^20^.

### 2.3 Dimensionality reduction algorithms and feature sets

Based on the dataset collected here, we proceeded to identify the feature importance to predict damage resulting from stimulation of neural tissue. For this, we first transformed categorical features into numerical ones using One-Hot encoding^44^, which converts each category into a unique binary vector of 0s and 1s. We used this method because these features did not have a natural ordered relationship and were of relatively low cardinality (e.g., 4 categories for feature Waveform shape: monophasic, biphasic symmetric, biphasic discharge, biphasic asymmetric)^44^. Then, we used a supervised machine learning random forest algorithm^45^ to identify feature importance by measuring how much each feature reduces or increases the accuracy across multiple decision trees. Then, for additional visualization, we tested three linear and non-linear dimensionality reduction algorithms^46^: principal component analysis (PCA), t-distributed stochastic neighbor embedding (t-SNE), and uniform manifold approximation and projection (UMAP).

### 2.4 Machine learning model training and testing

A Dell desktop computer (Intel Core i7-10700 CPU @ 2.90 GHz, Windows 10 enterprise 64-bit OS and 32 GB RAM) and a MacBook Air (3.2 GHz 8-Core Apple M1, macOS Ventura 13 and 16 GB RAM) were used to conduct the machine learning modeling. Scikit-Learn, an open-source Python machine learning library, was used for dimensionality reduction and data visualization (dimension of the embedded space: 2 or 3, all other parameters set as default). Additionally, Scikit-Learn was used to conduct all model training and testing procedures. Other Python libraries, including NumPy, Pandas, and Matplotlib, were also included for data analysis and presentation.

Next, the complete dataset (385 entries) was randomly split, based on uniform distributions, into training and testing subsets with a ratio of 80:20 (training:testing). Both datasets were well-balanced between the two labels (for training, label 0: 151 entries and label 1: 157 entries; for testing, label 0: 44 entries and label 1: 33 entries). Finally, since the means of numerical features varied significantly across variables, standardization was performed on both training and testing datasets by subtracting from mean and dividing by standard deviation.

For the current study, four supervised learning algorithms were investigated^47,48^: logistical regression, k-nearest neighbor, random forest, and multilayer perceptron (MLP). Logistical regression is a linear model that predicts the probability of a binary outcome based on input features. K-nearest neighbor assigns a label to a new instance based on the majority vote of its closest neighbors. Random forest combines multiple decision trees to produce a robust and accurate prediction. MLP is a type of artificial neural network that can learn non-linear and complex relationships between features and outcomes. Hyperparameters used for each algorithm can be found in **Supplementary Table 1**.

We then proceeded to create three separate subsets, each with a different set of features. First, the Shannon feature set included only the charge per phase and charge density features, as previously described for the Shannon equation. Second, a semi-complete feature set included features that are readily available in retrospective studies but are sometimes unpredictable in experimental case-control studies. The features excluded were parameters relating to total stimulation delivered during the study: days, hours, number of pulses, and accumulated charge. In addition, animal model and specific tissue region targeted for stimulation were excluded to enable the creation of a web-based application that is capable of generalizing predictions to the nervous system regardless of experimental design for most user cases. After exclusions, the semi-complete set included 25 features (10 categorical and 15 numerical). Third, a partial feature set included the most important features as revealed by the one-hot random forest algorithm and features required to calculate those variables (1 categorical and 12 numerical).

A modeling pipeline inspired from our previous study was adopted^47^. First, we screened all models with the One-Hot encoded/standardized training dataset using tenfold cross-validation and adopted two filtering conditions: (1) mean accuracy > 0.75, and (2) standard deviation of accuracy < 0.10. Next, we applied all candidate models on the testing dataset and adopted secondary filtering conditions as: (1) accuracy > 0.85, and (2) AUC (Area Under the ROC Curve) > 0.85.

Furthermore, using the random forest algorithm, we proceeded to train and test the best-performing classifier on the damage level (e.g., 0 – no damage; 1 – minimal damage; 2 – mild damage; 3 – moderate damage; and 4 – severe damage), rather than the binary outcome. We first performed tenfold cross-validation (number of trees from 1-100) and subsequently applied the trained models on the testing dataset. However, it should be emphasized that due to the increase in cardinality for the output labels, the number of instances for each specific label were even lower than those of binary labeling.

Finally, for comparison purposes, we additionally explored the use of ordinal encoding for categorical feature transformations instead of One-Hot encoding, which was subsequently split and standardized into training and testing using the same analytical pipeline as detailed for One-Hot encoding for the best-performing feature set.

### 4.5 Performance metrics

Performance of each of the different models, including the Shannon equation, was evaluated using threshold dependent and independent metrics, which include:

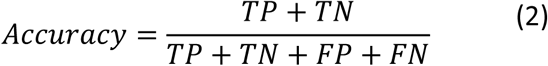

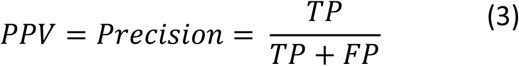

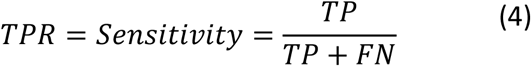

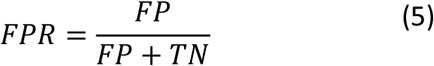

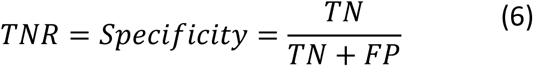

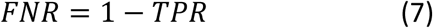

where TP denotes the number of entries correctly predicted as damaging (true positive), FP denotes the number of entries incorrectly predicted as damaging (false positive), TN denotes entries correctly predicted as non-damaging (true negative), and FN denotes entries incorrectly predicted as non-damaging (false negative). Accuracy (**Equation 2**) is a metric that calculates the success of correctly predicted cases of all entries. Precision or positive predictive value (PPV; **Equation 3**) measures the proportion of damaging cases that were actually damaging. The true positive rate (TPR; **Equation 4**), also known as recall or sensitivity, measures the model’s ability to correctly predict damaging cases from actual cases labeled as damaging. The false positive rate (FPR; **Equation 5**) measures the model’s capacity to incorrectly predict damaging cases given the actual cases labeled as non-damaging. The true negative rate (TNR; **Equation 6**) or specificity measures the proportion of non-damaging cases that were actually non-damaging. The false negative rate (FNR; Equation 7) measures the model’s capacity to incorrectly predict non-damaging cases given the actual cases labeled as damaging.

### 2.6 Statistical analysis

Descriptive statistics^49^ were calculated for all features and reported as mean ± SEM (Graphpad Prism 10, Version 10.0.2, Graphpad Software, LLC, Boston, MA, USA). First, a Shapiro-Wilk test was used to determine if features were normally distributed. Then, the non-parametric Kolmogorov-Smirnov test was used to reveal statistical differences between damaging and non-damaging parameters when data were not normally distributed. An unpaired t-test was used when data were normally distributed. To compare the sensitivity and specificity of each one of the models described here, including the Shannon equation, we used the McNemar test and exact binomial test in RStudio (2023.06.2, Posit Software, PBC, Boston, MA, USA) using the DTComPair package^50^.

### 2.7 Web hosting of machine learning algorithm

We designed a Java JSP (JavaServer Pages) based web application for implementing use of the best performing algorithm with the least possible input features. The web application was hosted at Amazon AWS (Amazon web services) using the EC2 t3a.micro instance (neurostimML.utdallas.edu).

## 3. Results

### 3.1 Dataset of electrical neurostimulation parameters and outcomes

The literature search for neurostimulation parameters and neural tissue outcomes yielded a total of 444 publications in PubMed, and an additional 42 relevant articles were found in literature reviews and books^20,21,39^. A total of 58 publications (references in **Supplementary Table 2**) were included in the database (**Supplementary Data 1**). From these publications, we extracted 385 neuromodulation measurements and their respective histological or functional responses. Responses were scored by the original authors for 60.3% (232) of the included entries; the remainder 39.7% (153) were scored by two independent investigators. There was a correlation of ρ=0.88 between the two investigators, which showed differences in only 36 entries. Binary scores were then calculated based on the averaged graded responses between investigators into damaging and non-damaging. This resulted in 50.7% (195) entries being labeled as non-damaging and 49.3% (190) as damaging, which demonstrated a balanced ratio of nearly 1:1, which is crucial for effective implementation of learning algorithms including Logistic Regression, K-nearest neighbor, Random Forest and Multilayer Perceptron^40,41^.

We graphically depicted the distribution of six commonly reported numerical features (**Supplementary Figure 2**) in the collection per graded tissue outcome, including electrode GSA, pulse width, frequency, current amplitude, charge per phase, and charge density. **Table 2** shows the median and range of the numerical features for the overall collection, and for damaging and non-damaging entries. In addition, it shows the p-values of the Kolmogorov-Smirnov test between both groups for all numerical features. This feature space represents the most comprehensive collection of neuromodulation parameters and tissue outcomes reported to date. There were 15 out of 19 (79%) numerical features that presented statistical differences between non-damaging and damaging parameters. Consistent with the Shannon equation, the charge per phase appeared to have a statistical significance (p=0.0001) between these two groups (damaging: 1.4±0.32 µC ph^-1^ vs. non-damaging: 0.27±0.05 µC ph^-1^); however, the charge density per phase did not reach statistical significance (damaging: 247.7±37.9 µC cm^-2^ ph^-1^ vs. non-damaging: 239.2±39.9 µC cm^-2^ ph^-1^; p=0.07). The mean of the damaging parameters had the following characteristics: larger GSA = 0.04±0.007 cm^2^ (vs. non-damaging: 0.03±0.006 cm^2^; p=0.0002), longer pulse width = 258.8±18.1 µs (vs. non-damaging: 195.0±16.4 µs; p=0.005), similar frequency = 267.1±32.5 Hz (vs. non-damaging: 330.9±38.8 Hz; p=0.07), and higher current = 3.2±0.5 mA (vs. non-damaging: 1.3±0.15 mA; p=0.008). These results suggest that using only the two variables of the Shannon equation: charge per phase and charge density, to predict stimulation-induced neural tissue damage is not sufficient, and other parameters such as frequency likely play a determinant role.

**Table 2.** Summary statistics of numerical features included in collection sorted by p-value of Kolmogorov-Smirnov test between damaging and non-damaging entries. Note that a value of 0.0 usually represents a value smaller than the number of significant digits.

| Feature | Overall<br>Median (Min–Max) | Units | Non-damaging<br>Median (Min–Max) | Damaging<br>Median (Min–Max) | p-value |  |
| --- | --- | --- | --- | --- | --- | --- |
| Total charge per phase accumulated | 3.6 (0.0–6830) | C | 0.4 (0.0–6831) | 8.5 (0.0–4479) | **** | <0.0001 |
| Charge per phase accumulated daily | 0.4 (0.0–22.7) | C | 0.1 (0.0–7.6) | 0.7 (0.0–23) | **** | <0.0001 |
| Stimulation on | 4 (0.0–50) | h | 1 (0.0–50) | 7 (0–50) | **** | <0.0001 |
| Total stimulation per day | 10.6 (8–24) | h | 7 (0.0–24) | 9 (0–24) | *** | 0.0001 |
| Charge per phase | 0.1 (0.0–48) | $\mu\text{C ph}^{-1}$ | 0.0 (0–6) | 0.1 (0.0–48) | *** | 0.0001 |
| GSA | 0.004 (0.0–0.50) | $\text{cm}^2$ | 0.002 (0.0–0.5) | 0.006 (0.0–0.5) | *** | 0.0002 |
| Number of pulses per day | 0.2 (0.0–22.5) | $\times 10^7$ pulses | 0.2 (0.0–22.5) | 0.2 (0.0–11.5) | *** | 0.0007 |
| Voltage amplitude | 1.0 (0.0–60) | V | 1.0 (0–19) | 1.0 (0.0–60) | ** | 0.002 |
| Total number of pulses | 0.009 (0.0–24.2) | $\times 10^9$ pulses | 0.01 (5–24.2) | 0.5 (0.0–21.6) | ** | 0.003 |
| Pulse width | 150 (25–1000) | $\mu\text{s}$ | 110 (25–975) | 200 (25–1000) | ** | 0.005 |
| Current density | 272.7 (0.0–32000) | $\text{mA cm}^{-2}$ | 363 (2.5–26084) | 273 (0.0–32000) | ** | 0.007 |
| Current amplitude | 1.0 (0.0–48) | mA | 1.0 (0–15) | 1 (0.0–48) | ** | 0.008 |
| Total stimulation | 36 (0.0–51840) | h | 36 (0.0–51840) | 36 (0.0–38880) | ** | 0.009 |
| Number of days stimulated | 7 (1–2160) | days | 18 (1–2160) | 4 (1–1620) | * | 0.04 |
| Duty cycle | 100 (0.0–100) | % | 100 (0–100) | 100 (1–100) | * | 0.04 |
| Frequency | 130 (1–2750) | Hz | 130 (1–2750) | 100 (1–2000) | ns | 0.07 |
| Charge density per phase | 39.2 (0.3–5216) | $\mu\text{C cm}^{-2} \text{ph}^{-1}$ | 34 (0.3–5217) | 71 (0.3–3200) | ns | 0.07 |
| Stimulation off | 0.0 (0.0–24) | h | 0.0 (0.0–24) | 0 (0–8) | ns | 0.45 |
| Interphase delay | 0.0 (0.0–800) | $\mu\text{s}$ | 0.0 (0–800) | 0.0 (0.0–400) | ns | >0.99 |

### 3.2 Performance assessment of the Shannon equation

**Figure 1A** shows a linear boundary widely accepted to be set at *k* = 1.85 (accepted values range between *k* = 1.5 and *k* = 2.0) that can separate the log-values of charge per phase and charge density for the dataset that gave rise to the Shannon equation. However, plotting the log-values for all entries included in this database (**Figure 1B**) revealed that the Shannon equation cannot linearly separate between damaging and non-damaging stimulation parameters. Even when attempting to predict tissue outcomes with the two additional linear boundaries (modified Shannon equation) that are frequently used in combination with the Shannon equation^21^ – prediction as non-damaging charge densities up to 30 µC cm^-2^ ph^-1^ for Medtronic macroelectrodes with GSA > 0.03 cm^2^ (regardless of *k*-value) and a maximum charge per phase of 4 nC ph^-1^ for microelectrodes with GSA < 2000 µm^2^ (regardless of *k*-value) – did not appear to accurately predict electrical stimulation-induced neural tissue outcomes. To fully assess the accuracy of the standalone Shannon equation and the modified Shannon equation, we first calculated the k-values for all entries in the collection. Then, we evaluated the accuracies to predict tissue outcomes using different k-values ranging from *k* = -4 to *k* = 4.

**Figure 1.**
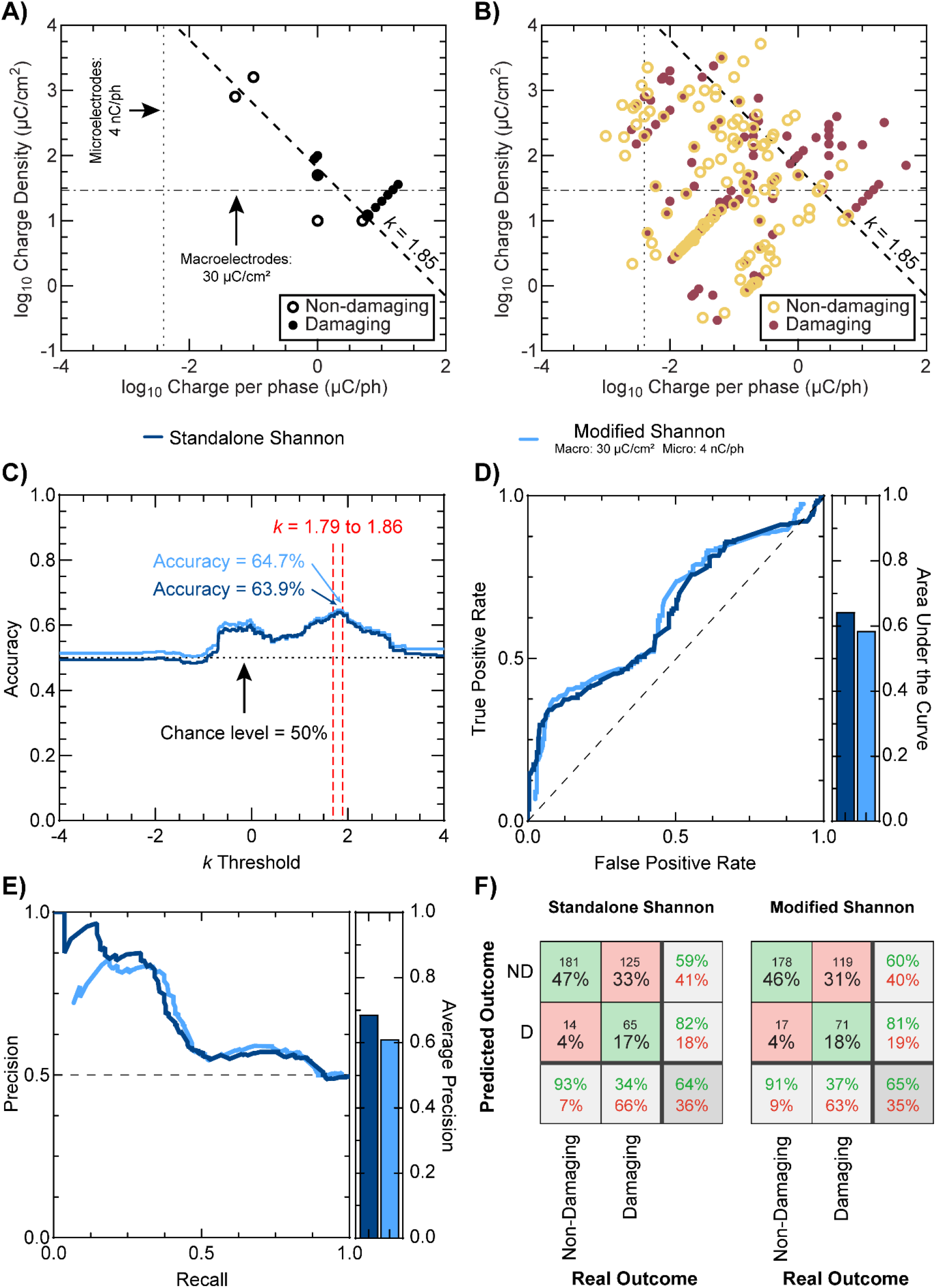
Prediction of electrical stimulation-induced neural tissue outcomes using the Shannon equation. **A)** Log-values of charge per phase and charge density per phase showing the dataset used for creation of the Shannon equation. **B)** Log-values of charge per phase and charge density per phase showing all data in this collection (including the Shannon equation dataset). **A&B)** show damaging and non-damaging regions of neuromodulation with a commonly used boundary of *k*=1.85 for the Shannon equation (dashed line), a charge density per phase for macroelectrodes commonly set at 30 µC/cm^2^ (dash-dotted line), and a maximum charge per phase for microelectrodes commonly set at 4 nC/ph (dotted line). Border only symbols depict non-damaging parameters; filled symbols depict damaging parameters. **C)** Accuracies achieved with the standalone Shannon equation (dark blue line) and the modified Shannon equation (30 µC/cm^2^ for macroelectrodes and 4 nC/ph for microelectrodes; light blue) for thresholds between *k*=0 and *k*=4. **D)** Receiver-operator curves (left) of predictions with the standalone Shannon and modified Shannon equations, and their respective areas under the curve (right). **E)** Precision-recall curve (left) for the standalone Shannon and modified Shannon equations, and their respective areas under the curve (right). **F)** Confusion matrices for prediction with the standalone Shannon equation (left) and the modified Shannon equation (right) using the best-performing threshold of *k*=1.85.

Consistent with previous observations^21^, the maximum accuracy (standalone Shannon: 63.9% and modified Shannon: 64.7%) was achieved with k-values ranging from k = 1.79 to k = 1.86 (**Figure 1C**); however, the use of the two boundaries in combination with the Shannon equation did not improve the accuracy. These values were found to be only 14% above the chance level (50%) indicating poor performance of both classifiers and suggesting that both classifiers may not fully capture the complex mechanisms of electrical stimulation-induced neural tissue damage. The performance of both, the standalone Shannon and modified Shannon equations was further investigated, as shown in **Figures 1C-F**, by looking at the receiver-operator (ROC) and precision-recall curves and confusion matrices.

Results show that the Shannon equation has low false positive rates (damaging outcome incorrectly predicted as non-damaging) but also low true positive rates (damaging outcome correctly predicted as damaging) and the area under the curve (AUC) was found to be only slightly above the chance level of 0.50 (standalone Shannon: 0.64 and modified Shannon: 0.59), indicating a poor ability to distinguish between damaging and non-damaging parameters. Furthermore, the precision-recall curves confirm the previous observations (standalone Shannon: 0.68 and modified Shannon: 0.59). Finally, the confusion matrices revealed that both Shannon classifiers using *k* = 1.85 predicted between 63% to 66% of parameters that were damaging as non-damaging, and between 7% to 9% of parameters that were non-damaging as damaging. This suggests that the Shannon equation has a tendency to underestimate potentially damaging stimulating parameters. This could be explained due to the limited feature set used by the Shannon equation (i.e., charge and charge density) that does not take into consideration other variables that may play a role as significant predictors (e.g., pulse width).

### 3.3 Feature selection and data visualization to improve prediction of electrical stimulation-induced tissue outcomes

To identify which features may be most critical for predicting damage in neural tissue, we used a random forest algorithm, where ordinal encoding was used to transform categorical features into numerical formats. **Table 3** shows all features sorted by order of importance score. Unsurprisingly, the five most important features were the waveform shape, daily accumulated charge, charge density per phase, charge per phase, and the total number of pulses delivered throughout the study. The five least important features were circuit control (whether stimulation was current- or voltage-controlled), DC bias voltage, interphase delay, charge transfer mechanism, and nervous system (central or peripheral) targeted for stimulation. Then we attempted to find a combination of two features that could more accurately distinguish between damaging and non-damaging stimulation parameters. **Figure 2A** shows the two most important numerical features in a log scale against each other (charge density vs. daily accumulated charge per phase). **Supplementary Figure 3** shows four additional combinations of the most important features. Results for all feature combinations revealed that there was no possible two-feature linear classifier that could accurately separate damaging and non-damaging parameters. Subsequently, we visualized the data in the collection after attempting three linear and non-linear dimensionality reduction methods (**Figures 2B-D**): principal component analysis (PCA), t-distributed stochastic neighbor embedding (t-SNE), and uniform manifold approximation and projection (UMAP). However, plots revealed overlapping features between damaging and non-damaging stimulating parameters when visualizing in a 2-dimensional space. Even when visualizing a 3-dimensional space for the principal components (**Supplementary Figure 4**), we did not observe a linearly separable feature space. Indeed, the first three principal components from PCA captured relatively low percentages of total variance (26.5%, 15.4% and 12.9%, respectively), implying the necessity of higher dimensional feature spaces. Because of this, we proceeded to train three machine learning approaches capable of non-linear classification: K-nearest Neighbor, Random Forest, and Multi-layer Perceptron.

**Figure 2.**
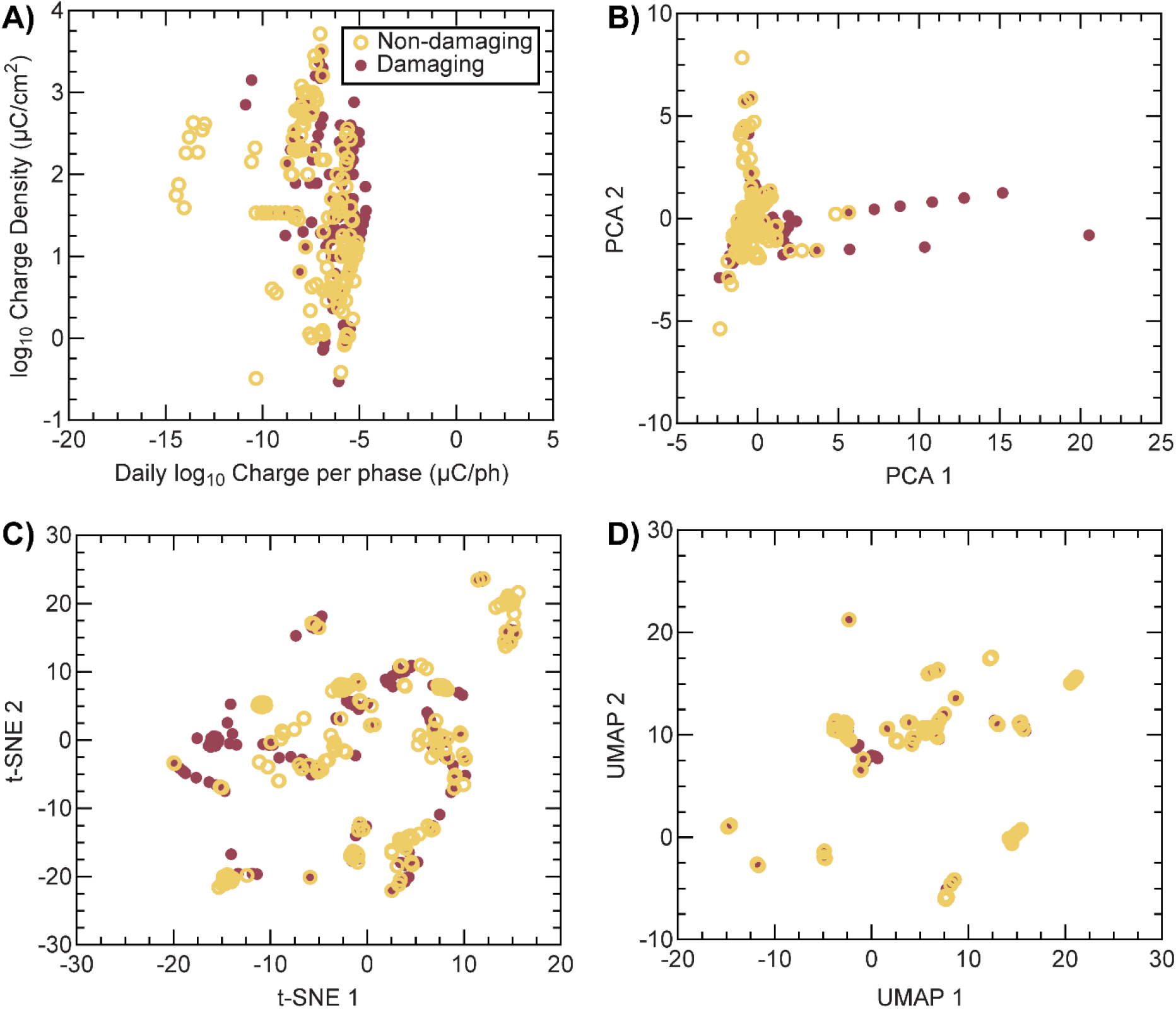
Feature selection and data visualization. Random forest was used to identify features which may be most critical for predicting damage in neural tissues. A threshold of 0.029 for importance score was used for feature selection. **A)** Log plot of two of the most important features (Charge Density vs. Daily Charge per phase). **B)** Visualization of the standardized dataset using PCA. **C)** Visualization of the standardized dataset using t-SNE. **D)** Visualization of the standardized dataset using UMAP. For all plots, border only symbols depict non-damaging parameters; filled symbols depict damaging parameters

**Table 3.** Feature importance score for accurate prediction of electrical stimulation-induced tissue outcomes.

| Importance | Feature | Score |
| --- | --- | --- |
| 1 | Waveform_shape | 0.07 |
| 2 | Daily_accumulated_charge | 0.05 |
| 3 | Charge_density | 0.05 |
| 4 | Charge_per_phase | 0.05 |
| 5 | Total_pulses | 0.05 |
| 6 | Tissue_target_structure | 0.04 |
| 7 | Total_accumulated_charge | 0.04 |
| 8 | Current_density | 0.04 |
| 9 | Daily_pulses | 0.04 |
| 10 | Stim_time_daily | 0.04 |
| 11 | Voltage_amplitude | 0.04 |
| 12 | Pulse_width | 0.04 |
| 13 | Days_stim | 0.04 |
| 14 | Animal | 0.04 |
| 15 | Current_amplitude | 0.04 |
| 16 | Stim_total_study | 0.03 |
| 17 | Frequency | 0.03 |
| 18 | Electrode_GSA | 0.03 |
| 19 | Stim_time_on | 0.03 |
| 20 | Tissue_target_structure | 0.03 |
| 21 | Duty_cycle | 0.03 |
| 22 | Electrode_material | 0.02 |
| 23 | Initial_current_direction | 0.02 |
| 24 | Stim_time_off | 0.02 |
| 25 | Electrode_shape | 0.02 |
| 26 | Array_configuration | 0.02 |
| 27 | Nervous_system | 0.02 |
| 28 | Charge_mechanism | 0.01 |
| 29 | Interphase_delay | 0.01 |
| 30 | Voltage_dc_bias | 0.01 |
| 31 | Circuit_control | 0.00 |

### 3.4 Machine learning approaches using One Hot Encoding and the Shannon feature set

Standardization of variables in preparation for machine learning training and testing was performed on all numerical features. Then, the training and testing subsets were created, yielding well-balanced sets between damaging and non-damaging labels (training 1:0.96 and testing 1:1.33, expressed as damaging:non-damaging). Links to the GitHub location of all training and testing datasets, valid machine learning models, performance metrics, and feature subsets can be found in **Supplementary Table 3**.

The first machine learning approach used the Shannon feature set with only two features, charge and charge density, to predict stimulation-induced neural tissue outcomes. None of the four algorithms tested (Logistical Regression, K-nearest Neighbor, Random Forest, and Multilayer Perceptron) yielded predictive models which satisfied the first set of filtering conditions during the training stage. However, it is worth noting that when applied to testing data, the best candidate model using the Random Forest algorithm (number of trees: 23, named as RF-Shannon-23) outperformed the modified Shannon equation, as it showed a higher area under the receiver-operator curve (**Figure 3A**, 0.83 vs 0.59), and similar average precision (**Figure 3B**, 0.69 vs. 0.61). In addition, compared to the modified Shannon equation (**Figure 3C**), this model was found to be more accurate (79% vs. 65%), with a higher ability to correctly predict damaging parameters (TPR = 88% vs. 37%) which resulted in a lower FNR (12% vs. 63%), and a higher TNR (89% vs. 60%). However, this classifier was less precise (71% vs 81%) with higher FPR (27% vs. 9%). The improvement on specificity and sensitivity as denoted by the TNR and TPR, respectively, were both found to be statistically significant (specificity: p = 0.005 and sensitivity: p = 0.0001). These results in combination with accuracy suggest a moderate improvement on the performance metrics compared to the Shannon equation, and while interesting, we proceeded to investigate the performance metrics of classifiers with the semi-complete and partial feature sets to try to identify a more accurate classifier.

**Figure 3.**
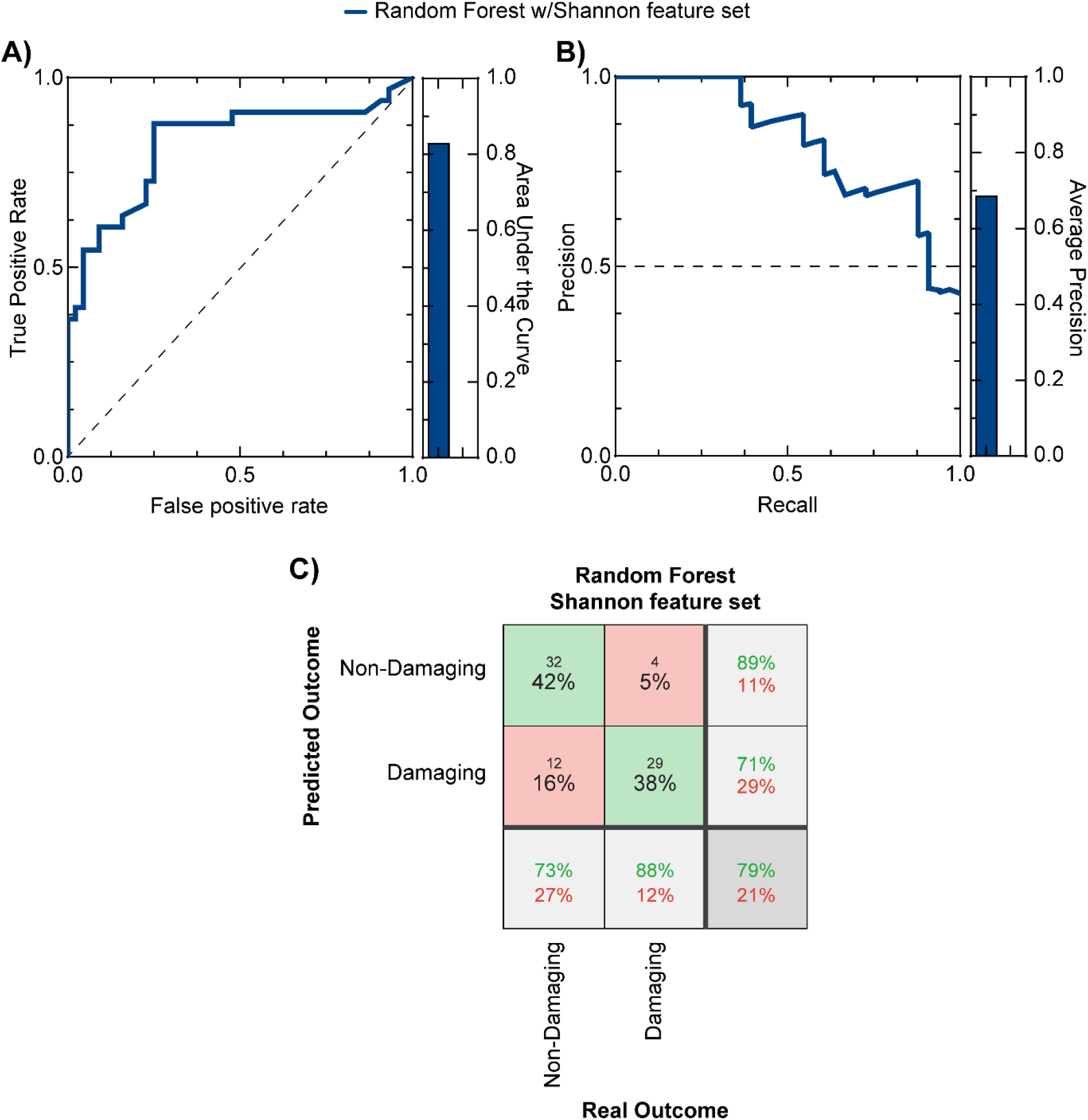
Predictive performance of the candidate machine learning approaches using the Shannon feature set and Random Forest algorithm. **A)** shows the Receiver-Operator curve (left) and the corresponding area under the curve (right) for the Random Forest (dark blue) algorithm. **B)** shows the precision-recall curve (left) and corresponding average precision (right) for the algorithm. Confusion matrix for the best-performing classifier that uses the Shannon feature set with random forest algorithm **C)** shows high accuracy (79%).

### 3.5 Machine learning approaches using One Hot Encoding and the semi-complete feature set

The second machine learning approach used the semi-complete feature set with 25 input features (**Supplementary Table 4**) that uses all features except for those that may not be readily available in experimental-control studies (e.g., total stimulation received). Similar to the Shannon feature set, neither Logistical Regression nor K-nearest Neighbor yielded valid predicting models. In contrast, the semi-complete feature set yielded 46 valid models using the Random Forest algorithm, and 445 valid models using the Multilayer Perceptron algorithm. Next, 14 Random Forest models and 46 Multilayer Perceptron models passed secondary filtering. Results for these models are shown in **Figure 4**, where the receiver operator curve with calculation of the area under the curve, and precision-recall curve with the average precision are shown. Results showed that the Random Forest had a lower area under the receiver operator curve (0.91 vs. 0.94) but higher precision (0.75 vs. 0.68) compared to the Multilayer Perceptron. The best-performing models for the Random Forest (number of trees: 89, named as RF-Total-89, accuracy = 87%, precision = 83%, TPR = 88%, FNR = 12%, TNR = 91%, FPR = 14%) and Multilayer Perceptron (number of nodes in first layer: 5, number of nodes in second layer: 26, named as MLP-Total-5-26, accuracy = 90%, precision = 86%, TPR = 91%, FNR = 9%, TNR = 93%, FPR = 11%) had better performance than both the modified Shannon equation and the Random Forest with the Shannon feature set. The specificity of both machine learning models was found to be not-statistically significant (Random Forest: p = 0.16 and Multilayer Perceptron: p = 0.32), whilst the sensitivity was found to be significantly improved for both models (Random Forest: p = 0.0001 and Multilayer Perceptron: p < 0.0001). These results support the use of the semi-complete feature set to more accurately predict damaging and non-damaging stimulation parameters. However, some features in this semi-complete set may be hard to estimate for certain stimulation protocols. Because of this, we aimed to further reduce the number of input features needed to predict stimulation-induced outcomes on neural tissue.

**Figure 4.**
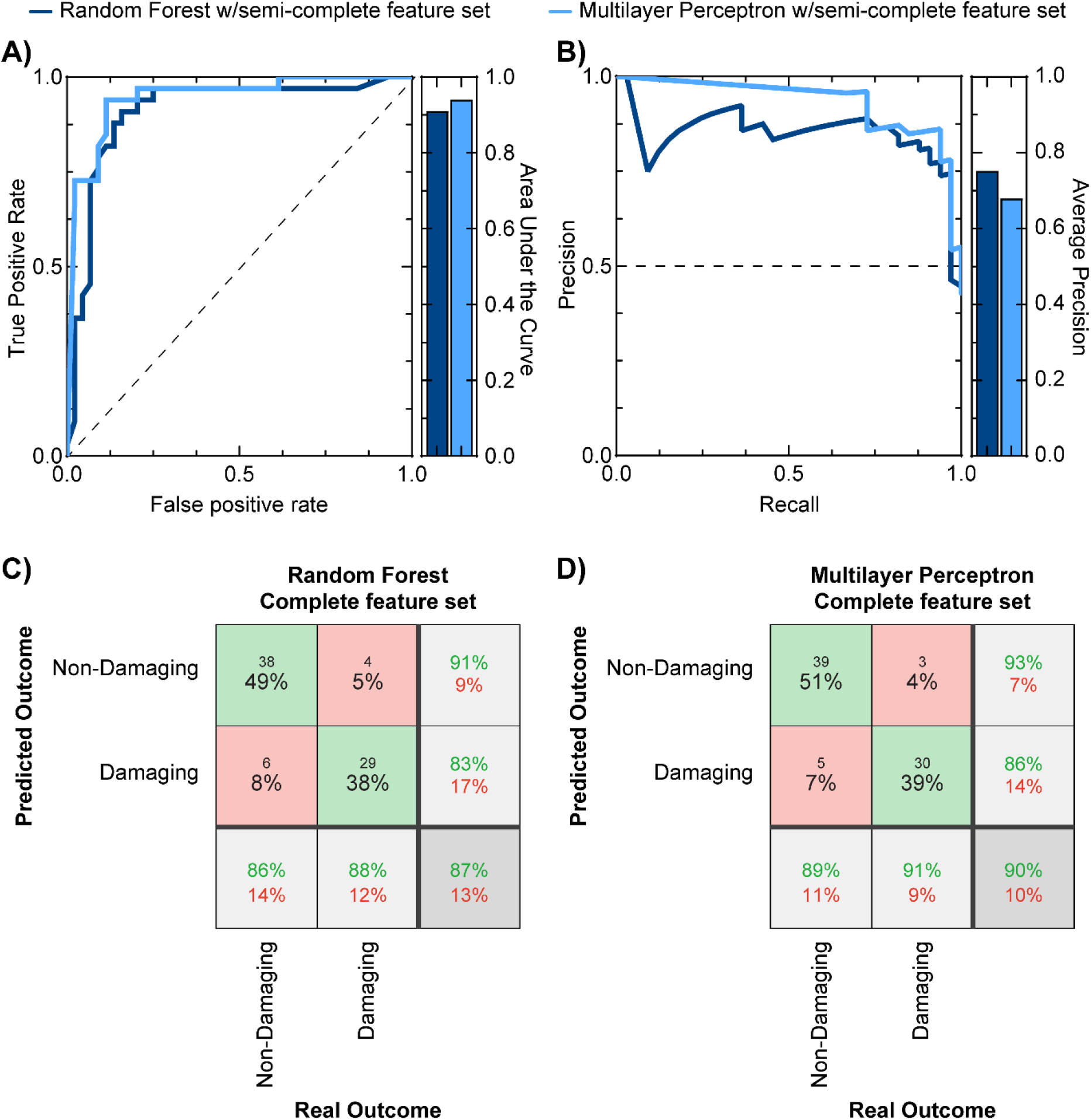
Predictive performance of two machine learning approaches using the semi-complete feature set. **A)** shows the Receiver-Operator curve (left) and the corresponding area under the curve (right) for both the Random Forest (dark blue) and the Multilayer Perceptron algorithms (light blue). **B)** shows the precision-recall curve (left) and corresponding average precision (right) for both algorithms. Confusion matrices for the best-performing classifiers that use the semi-complete feature set with random forest **C)** and multilayer perceptron **D)** algorithms show high accuracy (87% and 90% respectively) that was higher than the best-performing algorithm using the Shannon feature set (79% with the Random Forest algorithm).

### 3.6 Machine learning approaches using One Hot Encoding and the partial feature set

The third machine learning approach used a partial feature set with 13 input features (**Supplementary Table 5**). Logistic Regression and K-nearest neighbor did not yield any valid predicting models, so they were excluded from further analysis. We found 67 valid models for the partial feature set using the Random Forest algorithm and 720 valid models with the Multilayer Perceptron after the first round of filtering. After secondary filtering, we found 46 valid Random Forest models, and 9 valid Multilayer Perceptron models. **Figure 5** shows the performance of these two algorithms, where the Random Forest outperformed the Multilayer perceptron as evidenced by the area under the receiver operator curve (0.92 vs. 0.88, respectively) and the average precision (0.76 vs 0.65). The best-performing models for the Random Forest (number of trees: 19, named as RF-Partial-19, accuracy = 88%, precision = 80%, TPR = 97%, FNR = 3%, TNR = 97%, FPR = 18%) and Multilayer Perceptron (number of nodes in first layer: 10, number of nodes in second layer: 16, named as MLP-Partial-10-16, accuracy = 87%, precision = 81%, TPR = 91%, FNR = 9%, TNR = 93%, FPR = 16%) performed better than the modified Shannon equation and comparably to their counterparts using the semi-complete feature set. Compared to the modified Shannon equation, the specificity of the Random Forest model using the partial feature set was found to be statistically higher (p = 0.04) but the Multilayer Perceptron model with this feature set was not significant (p = 0.08). The sensitivity for both models was significantly higher than the modified Shannon equation (p < 0.0001). We then compared the models with the highest accuracies for the partial feature (Random Forest) set and the semi-complete feature set (Multilayer Perceptron) and found that both specificity (p = 0.08) and sensitivity (p = 0.16) were not statistically different. These results suggest that the best approach that maximizes accuracy, while minimizing user burden is the use of the partial feature set.

**Figure 5.**
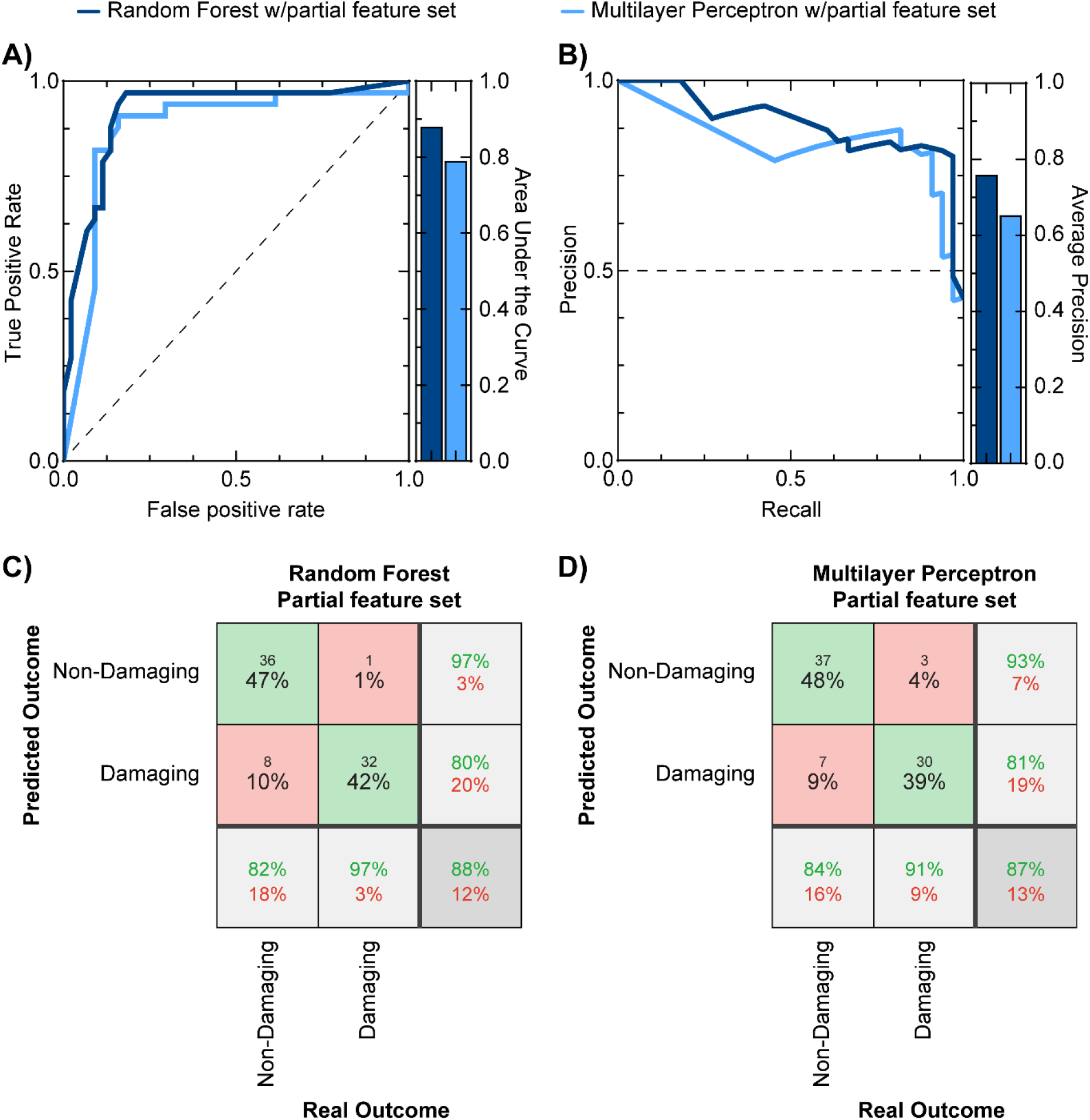
Predictive performance of two candidate machine learning approaches using the partial feature set. **A)** shows the Receiver-Operator curve (left) and the corresponding area under the curve (right) for both the Random Forest (dark blue) and the Multilayer Perceptron algorithms (light blue). **B)** shows the precision-recall curve (left) and corresponding average precision (right) for both algorithms. Confusion matrices for the best-performing classifiers that use the partial feature set with random forest **C)** and multilayer perceptron D) algorithms show high accuracy (88% and 87%, respectively) that was comparable to the best-performing algorithm with the semi-complete feature set (multilayer perceptron: 90%).

### 3.7 Machine learning approaches using One Hot Encoding and the partial feature set to predict non-binary damage level

Given that the Random Forest algorithm with the partial feature set had the best performance while minimizing the input features required, we proceeded to train a new algorithm to predict damage level (e.g., 0 – no damage; 1 – minimal damage, 2 – mild damage, 3 – moderate damage, and 4 – severe damage), rather than the binary outcome (1 – damaging, 0 – non-damaging). The training dataset had an unbalanced ratio with 33 instances labeled as level 0, 124 as level 1, 52 as level 2, 62 as level 3, and 37 as level 4. Similarly, the testing dataset had 8 labeled as level 0, 36 as level 1, 10 as level 2, 14 as level 3, and 9 as level 4. Because of the much smaller sample size, as expected, the overall accuracies were lower than those of the binary analysis. The maximum accuracy (number of trees: 8) achieved for this approach was 73% with an area under the receiver operator curve equal to 0.90 (**Supplementary Figure 5**). While the performance is better than the Shannon equation, the performance of the binary classifiers was superior and thus, we decided to proceed with the binary classifiers.

### 3.8 Machine learning approaches using ordinal encoding

As mentioned earlier, for constructing learning models, One-Hot encoding was used for categorical features. For comparison purposes, we additionally used ordinal encoding for categorical feature transformations, which was subsequently split and standardized into training (label 0: 155, label 1: 153) and testing (label 0: 40, label 1: 37) datasets. Next, we used the partial feature set which previously yielded high accuracy rates for the prediction of stimulation-induced outcomes on neural tissues. Only 9 models passed the first round of filtering conditions during tenfold cross-validation, compared to 67 valid models from One-Hot encoding. Next, when applied to the testing dataset, the best-performing model based on the Random Forest algorithm (number of trees: 13) yielded an accuracy of 81.8% and AUC of 0.86, both of which were lower than using One-Hot encoding (88.3% and 0.92, respectively). Taken together, these results showed a clear advantage of using One-Hot encoding over ordinal encoding for the categorical features in this study.

### 3.9 Web portal for Random Forest algorithm with partial feature set

We selected the Random Forest algorithm with the partial feature set to become publicly available on the web portal (neurostimML.utdallas.edu). A screenshot taken from the first version of the web portal can be found in **Figure 6**. The first screen is the submission of experimental condition values for the following parameters (**Figure 6A**, waveform shape, frequency, pulse width, current or voltage amplitude, electrode GSA, duty cycle, and stimulation duration) from the user, the values for all other features contained within the partial feature set are calculated from the input values and a prediction is made. For output, the predictive results from both our partial and the original Shannon equation are displayed (a sample output is shown in **Figure 6B**, likely no damage: green, likely tissue damage: magenta).

**Figure 6.**
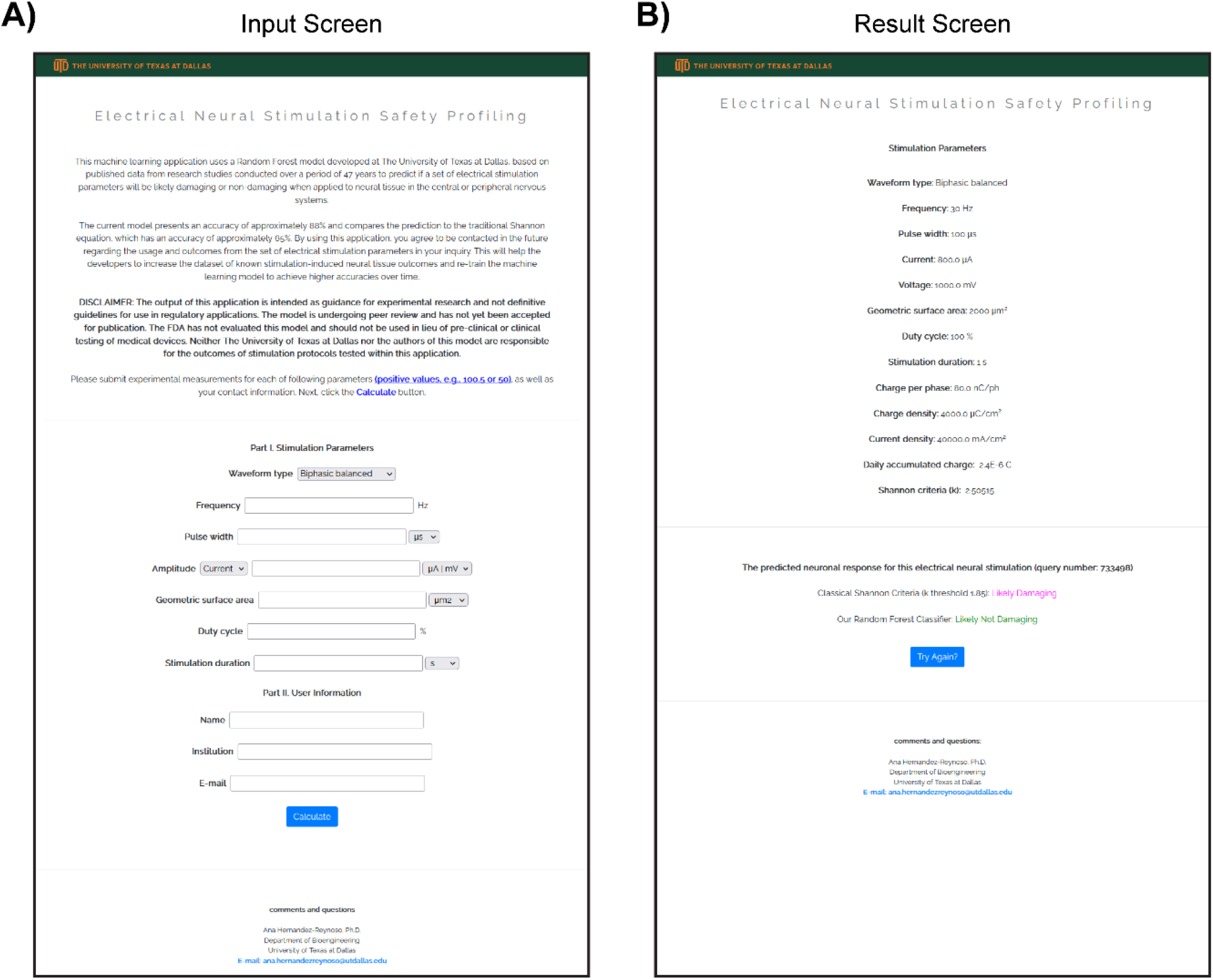
Web portal for Random Forest model using the partial feature set. **A)** A sample input page containing the parameter list consisting of waveform shape, frequency, pulse width, current or voltage amplitude, electrode GSA, duty cycle, and stimulation duration. **B)** A sample output page showing the predictive results from both our RF-Partial-19 and original Shannon equation (no damage: green, with damage: magenta).

## 4. Discussion

In the current study, we developed the most comprehensive dataset to-date for stimulation-induced neural tissue outcomes. Then, we developed a random forest-based machine learning classifier with a partial feature set which performed significantly better than the standalone and modified Shannon equations (accuracies: 88.3% and 64%, and 65% respectively). We have made this tool available online for use by the neural engineering field, as well as the complete literature dataset, to aid the generation of safer neural stimulation protocols.

The Shannon equation (standalone and modified) has been used and generally accepted since its inception more than 30 years ago. Here, we have demonstrated that it has a low ability to accurately predict tissue damage outcomes, especially when new electrode designs (e.g., microelectrodes), stimulation parameters (e.g., high frequency stimulation), and tissue regions are targeted for neuromodulation. The Shannon equation achieved a maximum accuracy of 64% to 65%, which is only moderately above the chance level of 50% accuracy. Most of the Shannon equation errors came from stimulation parameters that were predicted as non-damaging but were actually damaging (FNR between 63% to 66%). This is particularly concerning because such mislabeling may result in the clinical use of unsafe stimulation parameters. The use of machine learning approaches, even with the feature set employing only the Shannon input features, resulted in a significant improvement in prediction (FNR ranging between 3% to 12%); however, the partial feature sets yielded the best results (Random Forest: 3% and Multilayer Perceptron: 9%). These results suggest that the model made available for the public (Random Forest with Partial feature set, RF-Partial-19) and used for the construction of the web portal, does not tend to underestimate potentially damaging parameters, and does not significantly overestimate non-damaging parameters (FPR: 18%). Our findings suggest that our predictive model may offer a safer approach for stimulation parameter selection compared to that provided by the Shannon standalone and modified equations.

It should be noted that due to the nature of neuromodulation studies, our dataset displays two distinct properties: (1) it consisted of both categorical and numerical features; and (2) the dataset was relatively small with only 385 entries. The small dataset size was largely influenced by the fact that acquiring new relevant experimental results is both time-consuming and expensive. Consequently, for each of the proposed analyses, careful consideration was taken for the selection of testing and training subsets. Resulting subsets displayed the same probability distribution compared to the entire dataset by using random training-testing splitting and standardization (using means and standard deviations of numerical features of the training dataset), which is crucial to achieve generalization of the constructed model. Furthermore, it was interesting that for our dataset Random Forest performed slightly better than Multilayer Perceptron algorithms when using the Partial feature set (accuracies for best candidates were 88% and 87%, respectively). Two possible factors could be involved: (1) the nature of Random Forest, which is based on conditional decisions, allows better handling of a mixture of categorical and numerical features^42^; and (2) the relatively small dataset size of our study (total dataset: 385, and training dataset: 308) may be a more pronounced constraint for Multilayer Perceptron, which typically requires large datasets. As an example, using the Partial feature set (16 features), the number of parameters for a relatively simple two-layer MLP (MLP-Partial-10-16) will be 373 (for weights: 16*10 + 10*16 + 16*1 = 336, and for biases: 10 + 16 + 1 = 27), which is greater than the size of our training instances and potentially results in underfitting of the predictive models. Future studies will address the dataset size by expanding our literature search and by using data mining, web crawling algorithms, and other novel methods to automate feature extraction from published studies. However, an immediate challenge with dataset expansion is that published studies are not consistent in reporting stimulating parameters. To standardize reporting of stimulation parameters in neural engineering, we suggest that at least the following parameters are clearly reported in all publications: waveform type, frequency, pulse width, if stimulation is current- or voltage-controlled, pulse amplitude, geometric surface area of the electrodes, duty cycle (or train duration and time between stimulation trains), stimulation duration per day, number of days receiving stimulation, electrode shape, and electrode material.

Next, we noted that for evaluating electrical stimulation-induced tissue damage, a scaled output (e.g., 0 – no damage; 1 – minimal damage, 2 – mild damage, 3 – moderate damage, and 4 – severe damage) may provide more information compared to the binary labels used to generate models in this study. However, due to the small dataset size presented here, the accuracy was lower compared to that of the binary classifiers. It was worth noting that most of the errors encountered on the testing subset were only 1 level apart from the true outcome, indicating that if the dataset size is expanded, this approach has the potential to not only distinguish between damaging and non-damaging stimulation parameters, but also to indicate the level of expected damage.

A further limitation of this study was the lack of standardized methods for defining and reporting damaging and non-damaging stimulation parameters in neural tissue across literature. Here, we attempted to mitigate this limitation by having two blinded investigators assign levels based on reported descriptions. Our results suggested an agreement between both investigators, as highlighted by the high correlation between scores (0.88) that resulted in a difference in the binary outcome in less than 10% of entries. Future work can address this limitation by having more than two investigators score the outcomes, and by providing a standard definition of ‘damage’ for the neural engineering field to use in future reports. In addition, the field can benefit from consensus standards by reporting stimulation-induced neural tissue outcomes as reported here from 0 to 4 (0 – no damage; 1 – minimal damage, 2 – mild damage, 3 – moderate damage, and 4 – severe damage), which are consistent with previously reported outcomes by McCreery^11,12^ and Shepherd^18^.

Finally, we emphasize that incorporation of additional experimental data would allow further exploration of other modeling methods not tested here (e.g., support vector machines, deep learning) that require higher samples sizes than what was available from the literature search results generated in this study. Including data from parameter values not included in the current database may also lead to the identification of additional features that provide more insight into the relationship between specific stimulation protocols and specific types of tissue damage. In future work, we intend to utilize results from this machine learning based assessment to develop methods for targeted parameter searching and experimental design.

## 5. Conclusion

Despite the Shannon equation being used and generally accepted for over 30 years, recent investigations have challenged its utility, especially with novel neural interfaces and stimulation waveforms. Here we demonstrated that it has a limited accuracy in predicting stimulation-induced neural tissue outcomes. Furthermore, the dimension complexity of the feature space (e.g., charge per phase, frequency, pulse width etc.) does not allow for linear classifiers to accurately predict outcomes, even when non-linear transformations were applied. Machine learning algorithms, especially Random Forest and Multilayer Perceptron performed significantly better at predicting neural tissue outcomes, despite the limited size of the available dataset. We envision our model could provide more accurate predictions as more data is collected from existing published literature, the web portal, and from future studies and be a transformative tool for the development of safer or more reliable clinical electrical neuromodulation. Finally, as additional studies with novel stimulation protocols become available, the machine learning models may be retrained to adapt and robustly predict a larger range of stimulation parameters.

## Acknowledgements

Research reported in this publication was supported by the US National Science Foundation (NSF) grant 2114192 to L.B., a Cecil H. and Ida Green Endowment to L.B., The University of Texas at Dallas to L.B. and J.J.P., and the National Institute of Neurological Disorders and Stroke of the National Institutes of Health under Award Number K99NS135194 to A.G.H.-R. The content is solely the responsibility of the authors and does not necessarily represent the official views of the National Institutes of Health. The authors thank the work of Layan Dhaher for her assistance in the curation of stimulation parameters on an early version of the database. The authors also thank members from Dr. Leonidas Bleris’ lab, Dr. Taek Kang, John Nguyen, Zikun Zhou and Eleni Balla, for constructive discussions.

## Author Contributions

Conceptualization & Methodology: A.G.H.-R., R.F., and Y.L.; Formal Analysis: Y.L., R.F., D.G., and A.G.H.- R.; Software & Web Development: Y.L. and D.G.; Writing: all authors contributed to manuscript writing and editing; Supervision: S.F.C., J.J.P., L.B., and A.G.H.-R. Funding acquisition: J.J.P., L.B., and A.G.H.-R. All authors have read and agreed to the published version of the manuscript.

## Data Availability

All data collected for this article is available in the **Supplementary Materials**, and our GitHub depository: https://github.com/UTDyxl121030/BlerisLab/tree/Neuromodulation/

## References

1. Krauss, J. K. et al. Technology of deep brain stimulation: current status and future directions. Nature reviews. Neurology 17, 75–87 (2021).

2. Kay-Rivest, E., Schlacter, J. & Waltzman, S. B. Cochlear implantation outcomes in the older adult: a scoping review. Cochlear implants international 23, 280–290 (2022).

3. Toffa, D. H., Touma, L., El Meskine, T., Bouthillier, A. & Nguyen, D. K. Learnings from 30 years of reported efficacy and safety of vagus nerve stimulation (VNS) for epilepsy treatment: A critical review. Seizure 83, 104–123 (2020).

4. Caravaca, A. S. et al. An Effective Method for Acute Vagus Nerve Stimulation in Experimental Inflammation. Frontiers in neuroscience 13, 877 (2019).

5. Pruitt, D. T. et al. Usage of RePlay as a Take-Home System to Support High-Repetition Motor Rehabilitation After Neurological Injury. Games for health journal 12, 73–85 (2023).

6. Ganzer, P. D. et al. Closed-loop neuromodulation restores network connectivity and motor control after spinal cord injury. eLife 7, (2018).

7. Zomkowski, K. et al. The effectiveness of different electrical nerve stimulation protocols for treating adults with non-neurogenic overactive bladder: a systematic review and meta-analysis. International urogynecology journal 33, 1045–1058 (2022).

8. Huang, Y. & Koh, C. E. Sacral nerve stimulation for bowel dysfunction following low anterior resection: a systematic review and meta-analysis. Colorectal disease : the official journal of the Association of Coloproctology of Great Britain and Ireland 21, 1240–1248 (2019).

9. Hachmann, J. T. et al. Epidural spinal cord stimulation as an intervention for motor recovery after motor complete spinal cord injury. Journal of neurophysiology 126, 1843–1859 (2021).

10. Eckermann, J. M. et al. Systematic Literature Review of Spinal Cord Stimulation in Patients With Chronic Back Pain Without Prior Spine Surgery. Neuromodulation : journal of the International Neuromodulation Society (2021) doi:10.1111/ner.13519.

11. McCreery, D. B., Agnew, W. F., Yuen, T. G. & Bullara, L. A. Comparison of neural damage induced by electrical stimulation with faradaic and capacitor electrodes. Annals of biomedical engineering 16, 463–481 (1988).

12. McCreery, D. B., Agnew, W. F., Yuen, T. G. & Bullara, L. Charge density and charge per phase as cofactors in neural injury induced by electrical stimulation. IEEE transactions on bio-medical engineering 37, 996–1001 (1990).

13. Agnew, W. F., Yuen, T. G. & McCreery, D. B. Morphologic changes after prolonged electrical stimulation of the cat’s cortex at defined charge densities. Experimental neurology 79, 397–411 (1983).

14. Brown, W. J. et al. Tissue reactions to long-term electrical stimulation of the cerebellum in monkeys. Journal of neurosurgery 47, 366–379 (1977).

15. Pudenz, R. H., Bullara, L. A., Jacques, S. & Hambrecht, F. T. Electrical stimulation of the brain. III. The neural damage model. Surgical neurology 4, 389–400 (1975).

16. Yuen, T. G., Agnew, W. F., Bullara, L. A., Jacques, S. & McCreery, D. B. Histological evaluation of neural damage from electrical stimulation: considerations for the selection of parameters for clinical application. Neurosurgery 9, 292–299 (1981).

17. Gilman, S., Dauth, G. W., Tennyson, V. M., Kremzner, L. T. & Defendini, R. Morphological and biochemical effects of chronic cerebellar stimulation in monkey. Transactions of the American Neurological Association 100, 9–14 (1975).

18. Shepherd, R. K. et al. Chronic intracochlear electrical stimulation at high charge densities: reducing platinum dissolution. Journal of neural engineering 17, 56009 (2020).

19. Rosenfeld, J. V et al. Tissue response to a chronically implantable wireless intracortical visual prosthesis (Gennaris array). Journal of neural engineering 17, 46001 (2020).

20. Shannon, R. V. A model of safe levels for electrical stimulation. IEEE transactions on bio-medical engineering 39, 424–426 (1992).

21. Cogan, S. F., Ludwig, K. A., Welle, C. G. & Takmakov, P. Tissue damage thresholds during therapeutic electrical stimulation. Journal of neural engineering 13, 21001 (2016).

22. Vatsyayan, R. & Dayeh, S. A. A universal model of electrochemical safety limits in vivo for electrophysiological stimulation. Frontiers in neuroscience 16, 972252 (2022).

23. Sarmiento Varón, L., et al. The role of machine learning in health policies during the COVID-19 pandemic and in long COVID management. Frontiers in public health 11, 1140353 (2023).

24. Smeele, N. V. R., Chorus, C. G., Schermer, M. H. N. & de Bekker-Grob, E. W. Towards machine learning for moral choice analysis in health economics: A literature review and research agenda. Social science & medicine (1982) 326, 115910 (2023).

25. Burns, B. L., Rhoads, D. D. & Misra, A. The Use of Machine Learning for Image Analysis Artificial Intelligence in Clinical Microbiology. Journal of clinical microbiology e0233621 (2023) doi:10.1128/jcm.02336-21.

26. Chen, R. J. et al. Algorithmic fairness in artificial intelligence for medicine and healthcare. Nature biomedical engineering 7, 719–742 (2023).

27. Roman-Naranjo, P., Parra-Perez, A. M. & Lopez-Escamez, J. A. A systematic review on machine learning approaches in the diagnosis and prognosis of rare genetic diseases. Journal of biomedical informatics 143, 104429 (2023).

28. Lee, M. Recent Advances in Deep Learning for Protein-Protein Interaction Analysis: A Comprehensive Review. Molecules (Basel, Switzerland) 28, (2023).

29. Nyangoh Timoh, K., et al. A systematic review of annotation for surgical process model analysis in minimally invasive surgery based on video. Surgical endoscopy 37, 4298–4314 (2023).

30. Yu, Z. et al. Disrupted network communication predicts mild cognitive impairment in end-stage renal disease: an individualized machine learning study based on resting-state fMRI. Cerebral cortex (New York, N.Y. : 1991) (2023) doi:10.1093/cercor/bhad269.

31. Kim, J. et al. Quantification of identifying cognitive impairment using olfactory-stimulated functional near-infrared spectroscopy with machine learning: a post hoc analysis of a diagnostic trial and validation of an external additional trial. Alzheimer’s research & therapy 15, 127 (2023).

32. Savage, J. T., Ramirez, J., Risher, W. C., Irala, D. & Eroglu, C. SynBot: An open-source image analysis software for automated quantification of synapses. bioRxiv : the preprint server for biology Preprint at 10.1101/2023.06.26.546578 (2023).

33. Zhang, Z., Zhang, Z., Ji, J. & Liu, J. Amortization Transformer for Brain Effective Connectivity Estimation from fMRI Data. Brain sciences 13, (2023).

34. Ghazi, N., Aarabi, M. H. & Soltanian-Zadeh, H. Deep Learning Methods for Identification of White Matter Fiber Tracts: Review of State-of-the-Art and Future Prospective. Neuroinformatics (2023) doi:10.1007/s12021-023-09636-4.

35. Zhou, H. et al. Super-resolution Segmentation Network for Reconstruction of Packed Neurites. Neuroinformatics 20, 1155–1167 (2022).

36. Kappattanavar, A. M., Hecker, P., Moontaha, S., Steckhan, N. & Arnrich, B. Food Choices after Cognitive Load: An Affective Computing Approach. Sensors (Basel, Switzerland) 23, (2023).

37. Kenkel, W. Automated behavioral scoring: Do we even need humans? Annals of the New York Academy of Sciences (2023) doi:10.1111/nyas.15041.

38. Qureshi, M. S., Qureshi, M. B., Asghar, J., Alam, F. & Aljarbouh, A. Prediction and Analysis of Autism Spectrum Disorder Using Machine Learning Techniques. Journal of healthcare engineering 2023, 4853800 (2023).

39. Shepherd, R. K. & McCreery, D. B. Basis of electrical stimulation of the cochlea and the cochlear nucleus. Advances in oto-rhino-laryngology 64, 186–205 (2006).

40. Maw, M., Haw, S.-C. & Ho, C.-K. Utilizing data sampling techniques on algorithmic fairness for customer churn prediction with data imbalance problems. F1000Research 10, 988 (2021).

41. Zhao, Z. et al. Conventional machine learning and deep learning in Alzheimer’s disease diagnosis using neuroimaging: A review. Frontiers in computational neuroscience 17, 1038636 (2023).

42. de Abreu Fontes, J., et al. Combining wavelength importance ranking to the random forest classifier to analyze multiclass spectral data. Forensic science international 328, 110998 (2021).

43. Miller, A. L. Effects of chronic stimulation on auditory nerve survival in ototoxically deafened animals. Hearing research 151, 1–14 (2001).

44. Wang, L. et al. Identifying sex-specific risk architectures for predicting amyloid deposition using neural networks. NeuroImage 275, 120147 (2023).

45. Wallace, M. L. et al. Use and misuse of random forest variable importance metrics in medicine: demonstrations through incident stroke prediction. BMC medical research methodology 23, 144 (2023).

46. Mitchell-Heggs, R., Prado, S., Gava, G. P., Go, M. A. & Schultz, S. R. Neural manifold analysis of brain circuit dynamics in health and disease. Journal of computational neuroscience 51, 1–21 (2023).

47. Li, Y. et al. Machine learning-based approaches for identifying human blood cells harboring CRISPR-mediated fetal chromatin domain ablations. Scientific reports 12, 1481 (2022).

48. Li, Y., Nowak, C. M., Pham, U., Nguyen, K. & Bleris, L. Cell morphology-based machine learning models for human cell state classification. npj Systems Biology and Applications 7, 1–9 (2021).

49. Kim, N., Fischer, A. H., Dyring-Andersen, B., Rosner, B. & Okoye, G. A. Research Techniques Made Simple: Choosing Appropriate Statistical Methods for Clinical Research. The Journal of investigative dermatology 137, e173–e178 (2017).

50. Stock, C., Hielscher, T. & Discacciati, A. DTComPair: comparison of binary diagnostic tests in a paired study design. Preprint at https://cran.r-project.org/package=DTComPair (2023).

