## Supplementary Materials for "NeurostimML: A machine learning model for predicting neurostimulation-induced tissue damage"

### A machine learning based approach for safe electrical neural stimulation

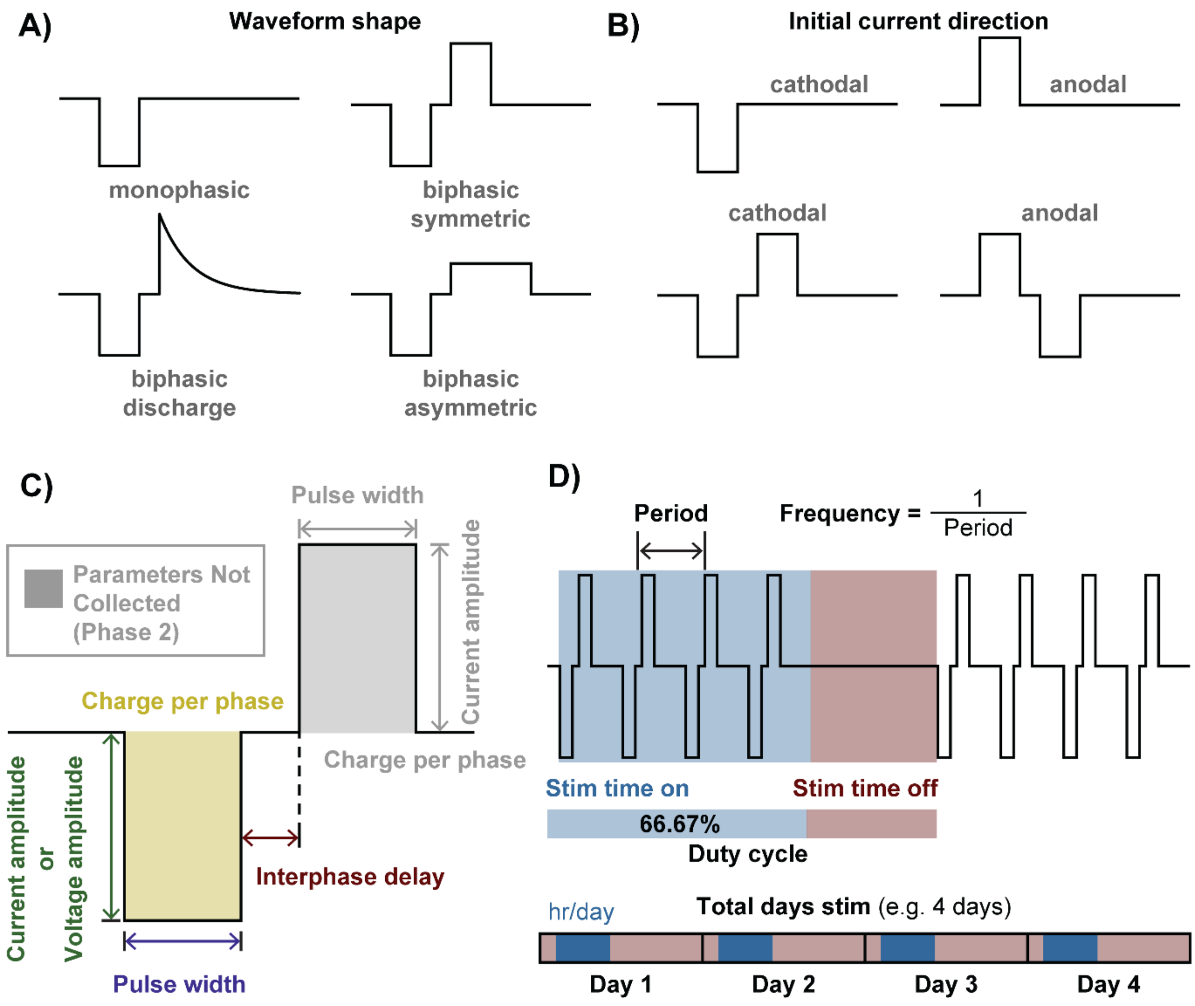


**Supplementary Figure 1. Visual definition of stimulation parameters and waveforms, includng A)** waveform shape, **B)** first phase direction, **C)** pulse parameters, **D)** frequency, duty cycle, and total days stimulated.

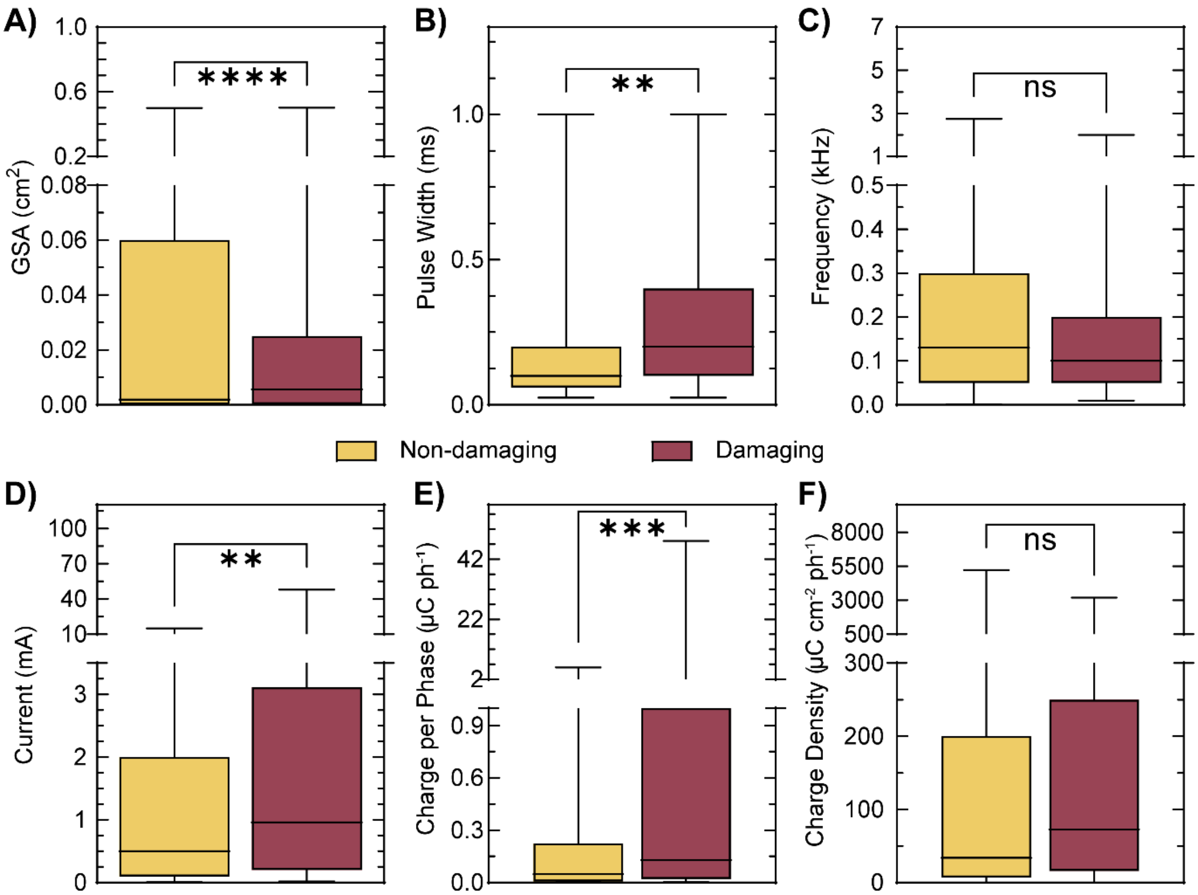


**Supplementary Figure 2. General statistics of selected features in this study.** Box plots showing the general statistics of selected features: **A)** GSA, **B)** Pulse Width, **C)** Frequency, **D)** Current, **E)** Charge per Phase, and **F)** Charge Density. For all plots, yellow: Non-damaging, red: Damaging.

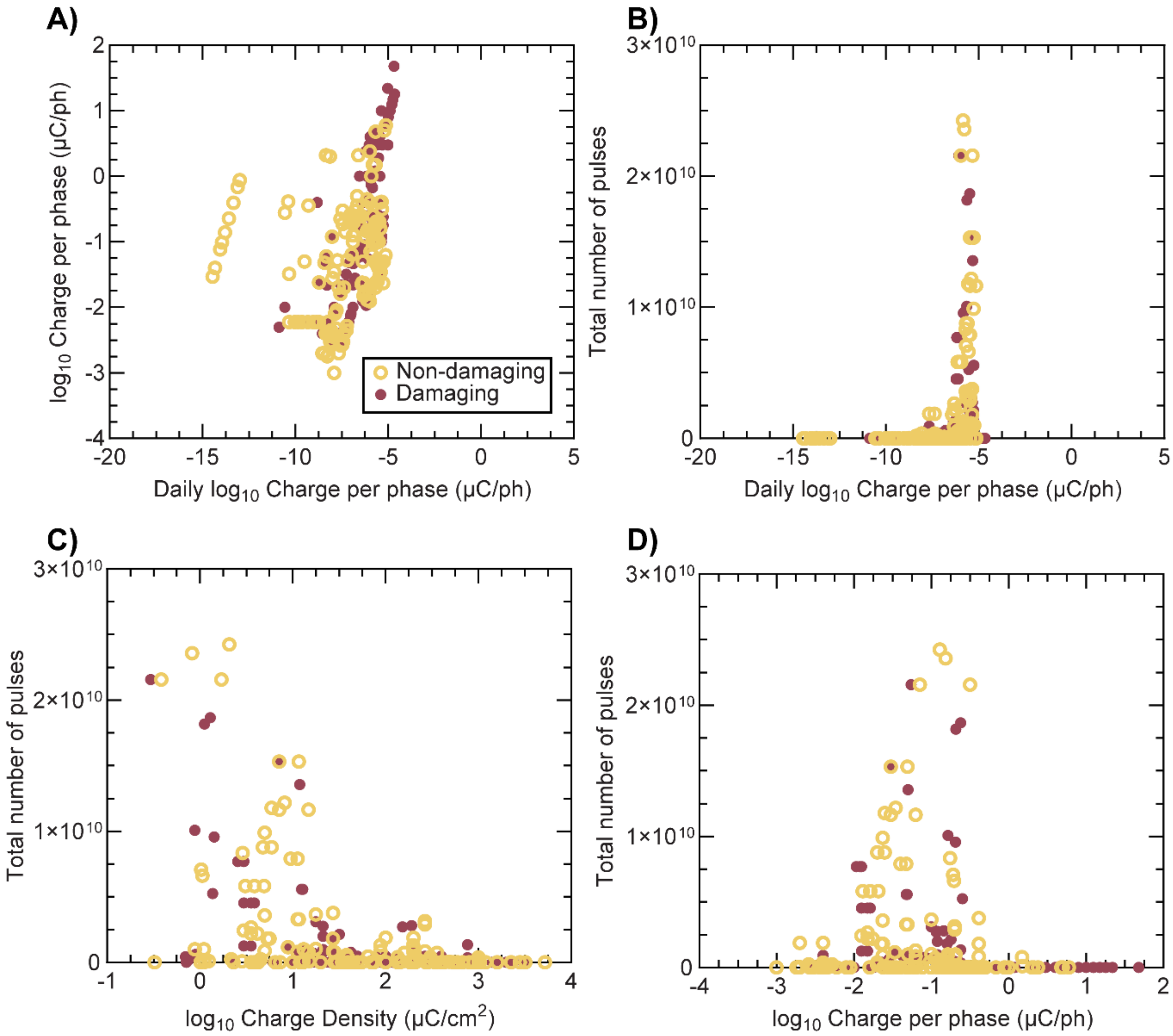


**Supplementary Figure 3. Feature selection and data visualization.** Random forest was used to identify features which may be most critical for predicting damage in neural tissues. A threshold of 0.029 for importance score was used for feature selection. Log plots show **A)** Charge per phase vs. Daily Charge per phase, **B)** Total number of pulses vs. Daily Charge per phase, **C)** Total number of pulses vs. Charge Density, and **D)** Total number of pulses vs. Charge per phase.

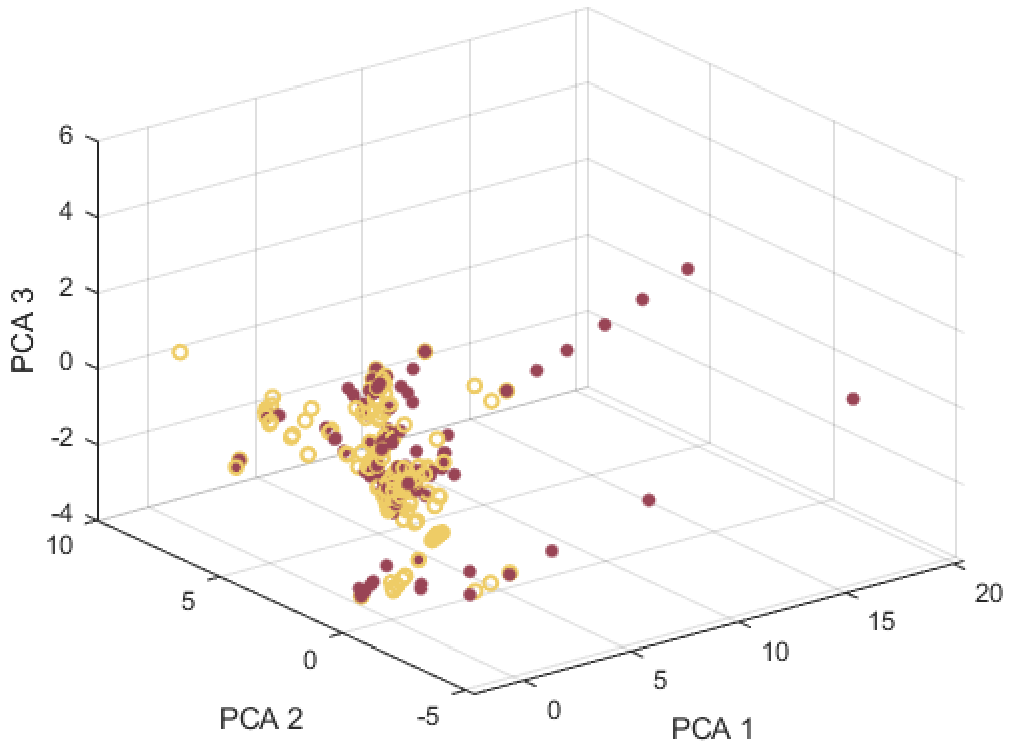


**Supplementary Figure 4. Feature selection and data visualization.** Visualization of the standardized dataset using the most important 3 principal components from PCA.

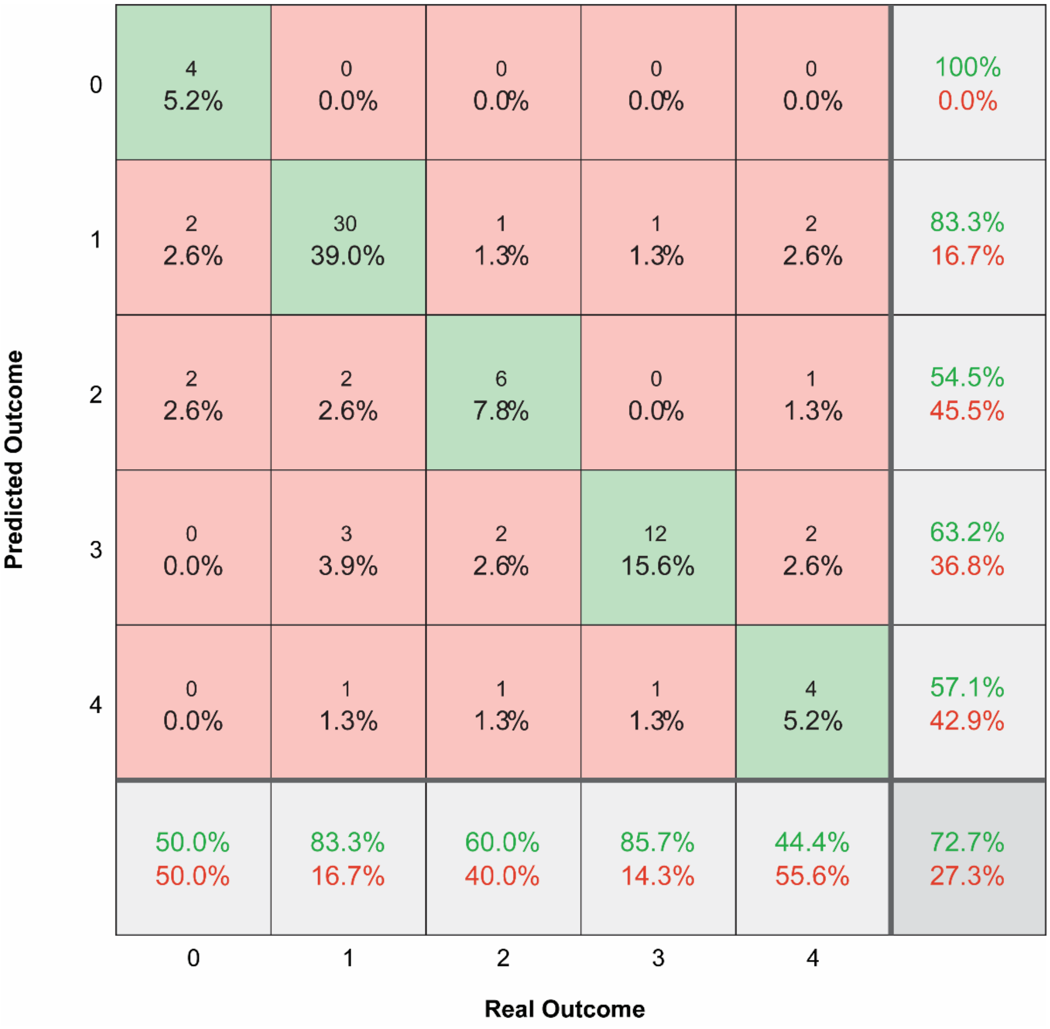


**Supplementary Figure 5. Multiclass confusion matrix for the best-performing Random Forest algorithm with partial feature set for the non-binary damage level**, where 0 – no damage; 1 – minimal damage, 2 – mild damage, 3 – moderate damage, and 4 – severe damage.

**Supplementary Table 1.** Hyperparameters used for training and testing machine learning algorithms.

| Machine Learning Algorithm | Hyperparameters |
|---|---|
| Logistic Regression | None |
| K-nearest neighbor (k-NN) | Number of trees: 1-100 |
| Random Forest (RF) | k: 1-100 |
| Multilayer Perceptron (MLP) | 2-hidden layers |
| | Nodes in first hidden layer: 1-30 |
| | Nodes in second hidden layer: 1-30 |
| | Activation function: tanh, relu, logistic |
| | Solver: adam, lbfgs, sgd |

**Supplementary Table 2.** References to studies included in the database.

| Reference ID | Reference |
|---|---|
| McCreery1990 | McCreery, D. B., Agnew, W. F., Yuen, T. G. H., & Bullara, L. (1990). Charge density and charge per phase as cofactors in neural injury induced by electrical stimulation. IEEE Transactions on Biomedical Engineering, 37(10), 996–1001. |
| McCreery1988 | McCreery, D. B., Agnew, W. F., Yuen, T. G. H., & Bullara, L. A. (1988). Comparison of neural damage induced by electrical stimulation with faradaic and capacitor electrodes. *Annals of Biomedical Engineering*, *16*(5), 463–481. |
| Agnew1983a | Agnew, W. F., Yuen, T. G. H., & McCreery, D. B. (1983). Morphologic changes after prolonged electrical stimulation of the cat's cortex at defined charge densities. *Experimental Neurology*, *79*(2), 397–411. |
| Brown1977 | Brown, W. J., Babb, T. L., Soper, H. V, Lieb, J. P., Ottino, C. A., & Crandall, P. H. (1977). Tissue reactions to long term electrical stimulation of the cerebellum in monkeys. *Journal of Neurosurgery*, *47*(3), 366–379. |
| Yuen1981 | Yuen, T. G. H., Agnew, W. F., Bullara, L. A., Jacques, S., & McCreery, D. B. (1981). Histological Evaluation of Neural Damage from Electrical StimulationConsiderations for the Selection of Parameters for Clinical Application. *Neurosurgery*, *9*(3), 292–299. |
| Yuen1984 | Yuen, T. G. H., Agnew, W. F., & Bullara, L. A. (1984). Histopathological Evaluation of Dog Sacral Nerve after Chronic Electrical Stimulation for Micturition. Neurosurgery, 14(4), 449–455. |
| Gilman1975 | Gilman, S., Dauth, G. W., Tennyson, V. M., Kremzner, L. T., & Defendini, R. (1975). Morphological and biochemical effects of chronic cerebellar stimulation in monkey. Transactions of the American Neurological Association, 100, 9–14. |
| Agnew1993 | Agnew, W. F., McCreery, D. B., Yuen, T. G. H., & Bullara, L. A. (1993). MK-801 protects against neuronal injury induced by electrical stimulation. *Neuroscience*, *52*(1), 45–53. |
| Pudenz1975 | Pudenz, R. H., Bullara, L. A., Jacques, S., & Hambrecht, F. T. (1975). Electrical stimulation of the brain. III. The neural damage model. *Surgical Neurology*, *4*(4), 389–400. |
| Schrader2006 | Schrader, L. M., Stern, J. M., Wilson, C. L., Fields, T. A., Salamon, N., Nuwer, M. R., Vespa, P. M., & Fried, I. (2006). Low frequency electrical stimulation through subdural electrodes in a case of refractory status epilepticus. *Clinical Neurophysiology*, *117*(4), 781–788. |
| Abejon2007 | Abejon, D., & Feler, C. A. (2007). Is impedance a parameter to be taken into account in spinal cord stimulation? *Pain Physician*, *10*(4), 533–540. |
| VNS1995 | George, R., Sonnen, A., Upton, A., Salinsky, M., Ristanovic, R., Bergen, D., Mirza, W., Rosenfeld, W., Nari-Toku, D., Manon-Espaillat, R., Barolat, G., Willis, J., Stefan, H., Treig, T., Hufnagel, A., Kuzniecky, R., Uthman, B., Wilder, B. J., Augustinsson, L., … Tarver, W. B. (1995). A randomized controlled trial of chronic vagus nerve stimulation for treatment of medically intractable seizures: The Vagus Nerve Stimulation Study Group*. *Neurology*, *45*(2), 224–230. |
| Kinoshita2004 | Kinoshita, M., Ikeda, A., Matsumoto, R., Begum, T., Usui, K., Yamamoto, J., Matsuhashi, M., Takayama, M., Mikuni, N., Takahashi, J., Miyamoto, S., & Shibasaki, H. (2004). Electric Stimulation on Human Cortex Suppresses Fast Cortical Activity and Epileptic Spikes. *Epilepsia*, *45*(7), 787–791. |

| | |
|---|---|
| Peyron2007 | Peyron, R., Faillenot, I., Mertens, P., Laurent, B., & Garcia-Larrea, L. (2007). Motor cortex stimulation in neuropathic pain. Correlations between analgesic effect and hemodynamic changes in the brain. A PET study. *NeuroImage*, *34*(1), 310–321. |
| Herzog2003 | Herzog, J., Volkmann, J., Krack, P., Kopper, F., Pötter, M., Lorenz, D., Steinbach, M., Klebe, S., Hamel, W., Schrader, B., Weinert, D., Müller, D., Mehdorn, H. M., & Deuschl, G. (2003). Two-year follow-up of subthalamic deep brain stimulation in Parkinson's disease. *Movement Disorders*, *18*(11), 1332–1337. |
| Caparros-Lefebvre1994 | Caparros-Lefebvre, D., Ruchoux, M., Blond, S., Petit, H., & Percheron, G. (1994). *Long-term t halamic stimulation Postmortem anatornoclinical study*. *October*. |
| Haberler2000 | Haberler, C., Alesch, F., Mazal, P. R., Pilz, P., Jellinger, K., Pinter, M. M., Hainfellner, J. A., & Budka, H. (2000). No Tissue Damage by Chronic Deep Brain Stimulation in Parkinson's Disease. |
| Burbaud2002 | Burbaud, P., Vital, A., Rougier, A., Bouillot, S., Guehl, D., Cuny, E., Ferrer, X., Lagueny, A., & Bioulac, B. (2002). Minimal tissue damage after stimulation of the motor thalamus in a case of chorea-acanthocytosis. Neurology, 59(12), 1982–1984. |
| DeBalthasar2008 | De Balthasar, C., Patel, S., Roy, A., Freda, R., Greenwald, S., Horsager, A., Mahadevappa, M., Yanai, D., McMahon, M. J., Humayun, M. S., Greenberg, R. J., Weiland, J. D., & Fine, I. (2008). Factors Affecting Perceptual Thresholds in Epiretinal Prostheses. Investigative Ophthalmology & Visual Science, 49(6), 2303–2314. |
| Mahadevappa2005 | Mahadevappa, M., Weiland, J. D., Yanai, D., Fine, I., Greenberg, R. J., & Humayun, M. S. (2005). Perceptual thresholds and electrode impedance in three retinal prosthesis subjects. IEEE Transactions on Neural Systems and Rehabilitation Engineering, 13(2), 201–206. |
| Veraart1998 | Veraart, C., Raftopoulos, C., Mortimer, J. T., Delbeke, J., Pins, D., Michaux, G., Vanlierde, A., Parrini, S., & Wanet-Defalque, M. C. (1998). Visual sensations produced by optic nerve stimulation using an implanted self-sizing spiral cuff electrode. Brain Research, 813(1), 181–186. |
| Fujikado2011 | Fujikado, T., Kamei, M., Sakaguchi, H., Kanda, H., Morimoto, T., Ikuno, Y., Nishida, K., Kishima, H., Maruo, T., Konoma, K., Ozawa, M., & Nishida, K. (2011). Testing of Semichronically Implanted Retinal Prosthesis by Suprachoroidal-Transretinal Stimulation in Patients with Retinitis Pigmentosa. Investigative Ophthalmology & Visual Science, 52(7), 4726–4733. |
| Xu1997 | Xu, J., Shepherd, R. K., Millard, R. E. & Clark, G. M. Chronic electrical stimulation of the auditory nerve at high stimulus rates: A physiological and histopathological study. Hear. Res. 105, 1–29 (1997). |
| McCreery1994 | McCreery, D. B., Yuen, T. G. H., Agnew, W. F., & Bullara, L. A. (1994). Stimulus parameters affecting tissue injury during microstimulation in the cochlear nucleus of the cat. Hearing Research, 77(1–2), 105–115. |
| McCreery1992 | McCreery, D. B., Yuen, T. G. H., Agnew, W. F., & Bullara, L. A. (1992). Stimulation with chronically implanted microelectrodes in the cochlear nucleus of the cat: Histologic and physiologic effects. *Hearing Research*, *62*(1), 42–56. |
| McCreery2000 | McCreery, D. B., Yuen, T. G. H., & Bullara, L. A. (2000). Chronic microstimulation in the feline ventral cochlear nucleus: physiologic and histologic effects. *Hearing Research*, *149*(1–2), 223–238. |

| | |
|---|---|
| McCreery2010 | McCreery, D., Pikov, V., & Troyk, P. R. (2010). Neuronal loss due to prolonged controlled-current stimulation with chronically implanted microelectrodes in the cat cerebral cortex. *Journal of Neural Engineering*, *7*(3), 036005. |
| McCreery2006 | McCreery, D., Lossinsky, A., Pikov, V., & Liu, X. (2006). Microelectrode array for chronic deep-brain microstimulation and recording. *IEEE Transactions on Biomedical Engineering*, *53*(4), 726–737. |
| McCreery1997 | McCreery, D. B., Yuen, T. G. H., Agnew, W. F., & Bullara, L. A. (1997). A characterization of the effects on neuronal excitability due to prolonged microstimulation with chronically implanted microelectrodes. IEEE Transactions on Biomedical Engineering, 44(10), 931–939. |
| McCreery1986 | McCreery, D. B., Bullara, L. A., & Agnew, W. F. (1986). Neuronal activity evoked by chronically implanted intracortical microelectrodes. Experimental Neurology, 92(1), 147–161. |
| Agnew1989 | Agnew, W. F., McCreery, D. B., Yuen, T. G. H., & Bullara, L. A. (1989). Histologic and physiologic evaluation of electrically stimulated peripheral nerve: Considerations for the selection of parameters. Annals of Biomedical Engineering, 17(1), 39–60. |
| Agnew1990b | Agnew, W. F., McCreery, D. B., Yuen, T. G. H., & Bullara, L. A. (1990). Local anaesthetic block protects against electrically-induced damage in peripheral nerve. Journal of Biomedical Engineering, 12(4), 301–308. |
| Kim1983 | Kim, J. H. et al. Light and electron microscopic studies of phrenic nerves after long-term electrical stimulation. J. Neurosurg. 58, 84–91 (1983). |
| Araki1998 | Araki, S. *et al.* Effects of chronic electrical stimulation on spiral ganglion neuron survival and size in deafened kittens. *Laryngoscope* **108**, 687–695 (1998). |
| Shepherd1994 | Shepherd, R. K., Matsushima, J., Martin, R. L. & Clark, G. M. Cochlear pathology following chronic electrical stimulation of the auditory nerve: II deafened kittens. Hear. Res. 81, 150–166 (1994). |
| Ni1992 | Ni, D. et al. Cochlear pathology following chronic electrical stimulation of the auditory nerve. I: Normal hearing kittens. Hear. Res. 62, 63–81 (1992). |
| Shepherd1983 | Shepherd, R. K., Clark, G. M. & Black, R. C. Chronic electrical stimulation of the auditory nerve in cats: Physiological and histopathological results. Acta Otolaryngol. 95, 19–31 (1983). |
| Walsh1982 | Walsh, S. M. & Leake-Jones, P. A. Chronic electrical stimulation of auditory nerve in cat: Physiological and histological results. Hear. Res. 7, 281–304 (1982). |
| Shepherd1999 | Shepherd, R. K., Linahan, N., Xu, J., Clark, G. M. & Araki, S. Chronic electrical stimulation of the auditory nerve using non-charge- balanced stimuli. Acta Otolaryngol. 119, 674–684 (1999). |
| Shepherd1991 | Shepherd, R. K., Matsushima, J., Millard, R. E. & Clark, G. M. Cochlear pathology following chronic electrical stimulation using non charge balanced stimuli. Acta Otolaryngol. 111, 848–860 (1991). |
| Mitchell1997 | Mitchell, A. et al. Effects of chronic high-rate electrical stimulation on the cochlea and eighth nerve in the deafened guinea pig. Hear. Res. 105, 30–43 (1997). |
| Agnew1986 | Agnew, W. F., Yuen, T. G. H., McCreery, D. B & Bullara, L. A. Histopathologic evaluation of prolonged intracortical electrical stimulation. Exp. Neurol. 92, 162–185 (1986). |
| McCreery1995 | McCreery, D. B., Agnew, W. F., Yuen, T. G. H & Bullara, L. A. Relationship between stimulus amplitude, stimulus frequency and neural damage during electrical stimulation of sciatic nerve of cat. Med. Biol. Eng. Comput. 33, 426–429 (1995). |

| | |
|---|---|
| Shepherd2021 | Shepherd, R. K. et al. Platinum dissolution and tissue response following long-term electrical stimulation at high charge densities. J. Neural Eng. 18, (2021). |
| Dalrymple2020 | Dalrymple, A. N. et al. Electrochemical and biological performance of chronically stimulated conductive hydrogel electrodes. J. Neural Eng. 17, (2020). |
| Rosenfeld2020 | Rosenfeld, J. V. et al. Tissue response to a chronically implantable wireless intracortical visual prosthesis (Gennaris array). J. Neural Eng. 17, (2020). |
| Shepherd2019 | Shepherd, R. K., Carter, P. M., Enke, Y. L., Wise, A. K. & Fallon, J. B. Chronic intracochlear electrical stimulation at high charge densities results in platinum dissolution but not neural loss or functional changes in vivo. J. Neural Eng. 16, (2019). |
| Bamford2017 | Bamford, J. A., Marc Lebel, R., Parseyan, K. & Mushahwar, V. K. The fabrication, implantation, and stability of Intraspinal Microwire arrays in the spinal cord of cat and rat. IEEE Trans. Neural Syst. Rehabil. Eng. 25, 287–296 (2017). |
| Orlowski2017 | Orlowski, D. et al. Brain Tissue Reaction to Deep Brain Stimulation—A Longitudinal Study of DBS in the Goettingen Minipig. Neuromodulation 20, 417–423 (2017). |
| Shepherd2017 | Shepherd, R. K., Wise, A. K., Enke, Y. L., Carter, P. M. & Fallon, J. B. Evaluation of focused multipolar stimulation for cochlear implants: A precli |
| Rajan2015 | Rajan, A. T. et al. The effects of chronic intracortical microstimulation on neural tissue and fine motor behavior. J. Neural Eng. 12, 066018 (2015). |
| Urdaneta2022 | Urdaneta, M. E. et al. The Long-Term Stability of Intracortical Microstimulation and the Foreign Body Response Are Layer Dependent. 16, 1–13 (2022). |
| Bamford2010 | Bamford, J. A., Todd, K. G. & Mushahwar, V. K. The effects of intraspinal microstimulation on spinal cord tissue in the rat. Biomaterials 31, 5552–5563 (2010). |
| Piallat2009 | Piallat, B. et al. Monophasic but not biphasic pulses induce brain tissue damage during monopolar high-frequency deep brain stimulation. Neurosurgery 64, 156–162 (2009). |
| Harnack2008 | Harnack, D. et al. Continuous high-frequency stimulation in freely moving rats: Development of an implantable microstimulation system. J. Neurosci. Methods 167, 278–291 (2008). |
| Manrique2008 | Manrique, M., Cervera-Paz, F. J., Insausti, A. M. & Nevison, B. Effects of cochlear nuclei electrical stimulation with surface brain stem implants in nonhuman primates. Ann. Otol. Rhinol. Laryngol. 117, 212–220 (2008). |
| Sun2008 | Sun, D. A. et al. Postmortem analysis following 71 months of deep brain stimulation of the subthalamic nucleus for Parkinson disease: Case report. J. Neurosurg. 109, 325–329 (2008). |
| Harnack2004 | Harnack, D. et al. The effects of electrode material, charge density and stimulation duration on the safety of high-frequency stimulation of the subthalamic nucleus in rats. J. Neurosci. Methods 138, 207–216 (2004). |

**Supplementary Table 3.** Links to the GitHub location of all training and testing datasets, valid machine learning models, performance metrics, and feature subsets.

| Filename | Description | GitHub Link |
|---|---|---|
| Supplementary Data 1 | Complete database with binary labels. | https://github.com/UTDyxl121030/BlerisLab/blob/Neuromodulation/Supplementary_Data1.xlsx |
| Supplementary Data 2 | Training subset after conversion of categorical features into numerical using One Hot encoding. | https://github.com/UTDyxl121030/BlerisLab/blob/Neuromodulation/Supplementary_Data2.xlsx |
| Supplementary Data 3 | Testing subset after conversion of categorical features into numerical using One Hot encoding. | https://github.com/UTDyxl121030/BlerisLab/blob/Neuromodulation/Supplementary_Data3.xlsx |
| Supplementary Data 4 | Performance metrics using Random Forest models and the semi-complete feature set. | https://github.com/UTDyxl121030/BlerisLab/blob/Neuromodulation/Supplementary_Data4.xlsx |
| Supplementary Data 5 | Performance metrics using Multilayer Perceptron models and the semi-complete feature set. | https://github.com/UTDyxl121030/BlerisLab/blob/Neuromodulation/Supplementary_Data5.xlsx |
| Supplementary Data 6 | Performance metrics using Random Forest models and the partial feature set. | https://github.com/UTDyxl121030/BlerisLab/blob/Neuromodulation/Supplementary_Data6.xlsx |
| Supplementary Data 7 | Performance metrics using Multilayer Perceptron models and the partial feature set. | https://github.com/UTDyxl121030/BlerisLab/blob/Neuromodulation/Supplementary_Data7.xlsx |
| Supplementary Data 8 | Performance metrics using Random Forest models, the partial feature set and non-binary labels. | https://github.com/UTDyxl121030/BlerisLab/blob/Neuromodulation/Supplementary_Data8.xlsx |
| Supplementary Data 9 | Training subset after conversion of categorical features into numerical using Ordinal encoding. | https://github.com/UTDyxl121030/BlerisLab/blob/Neuromodulation/Supplementary_Data9.xlsx |
| Supplementary Data 10 | Testing subset after conversion of categorical | https://github.com/UTDyxl121030/BlerisLab/blob/Neuromodulation/Supplementary_Data10.xlsx |

| | | |
|---|---|---|
| | features into numerical using Ordinal encoding. | |
| Supplementary Data 11 | Performance metrics using Ordinal encoding, Random Forest models and the partial feature set. | https://github.com/UTDyxl121030/BlerisLab/blob/Neuromodulation/Supplementary_Data11.xlsx |
| Supplementary Scripts | The RF-Partial-19 model, and other main python scripts used in this study. | https://github.com/UTDyxl121030/BlerisLab/blob/Neuromodulation/Supplementary_Scripts |

**Supplementary Table 4.** List of input features machine learning approaches utilizing the semi-complete feature set.

| Categorical | Numerical | |
|---|---|---|
| Nervous_system | Electrode_GSA | Current_density |
| Electrode_shape | Pulse_width | Frequency |
| Tissue_target_structure | Interphase_delay | Current_amplitude |
| Electrode_material | Voltage_amplitude | Charge_per_phase |
| Charge_mechanism | Charge_density | Duty_cycle |
| Waveform_shape | Stim_time_on | Stim_time_off |
| Initial_current_direction | Stim_time_daily | Daily_pulses_count |
| Array_configuration | Daily_accumulated_charge | |
| Circuit_control | | |
| Voltage_dc_bias | | |

**Supplementary Table 5.** List of input features machine learning approaches utilizing the partial feature set.

| Categorical | Numerical | |
|---|---|---|
| Waveform_shape | Electrode_GSA | Current_density |
| | Pulse_width | Stim_time_on |
| | Frequency | Stim_time_daily |
| | Current_amplitude | Daily_pulses_count |
| | Voltage_amplitude | Daily_accumulated_charge |
| | Charge_per_phase | |
| | Charge_density | |
